# A library of reporters of the global regulators of gene expression of *Escherichia coli*

**DOI:** 10.1101/2023.11.28.568972

**Authors:** Suchintak Dash, Rahul Jagadeesan, Ines S.C. Baptista, Vatsala Chauhan, Vinodh Kandavalli, Samuel M.D. Oliveira, Andre S. Ribeiro

**Affiliations:** Faculty of Medicine and Health Technology, Tampere University, 33520, Finland; Department of Cell and Molecular Biology, Uppsala University, Uppsala, Sweden; Joint School of Nanoscience and Nanoengineering, North Carolina A&T State University, Greensboro, 27401-NC, USA

**Keywords:** Global Regulators of Gene Expression, Transcriptional Reporters, Synthetic Single-Copy Plasmids, Native Promoters, Fluorescent Proteins, Strain Library

## Abstract

The topology of the transcription factor network (TFN) of *E. coli* is far from uniform, with 22 global regulator (GR) proteins controlling one-third of all genes. So far, their production rates cannot be tracked by comparable fluorescent proteins. We developed a library of fluorescent reporters for 16 GRs for this purpose. Each consists of a single-copy plasmid coding for GFP fused to the full-length copy of the native promoter. We tracked their activity in exponential and stationary growth, as well as under weak and strong stresses. We show that the reporters have high sensitivity and specificity to all stresses tested and detect single-cell variability in transcription rates. Given the influence of GRs on the TFN, we expect that the new library will contribute to dissecting global transcriptional stress-response programs of *E. coli*. Moreover, the library can be invaluable in bioindustrial applications that tune those programs to, instead of cell growth, favor productivity while reducing energy consumption.

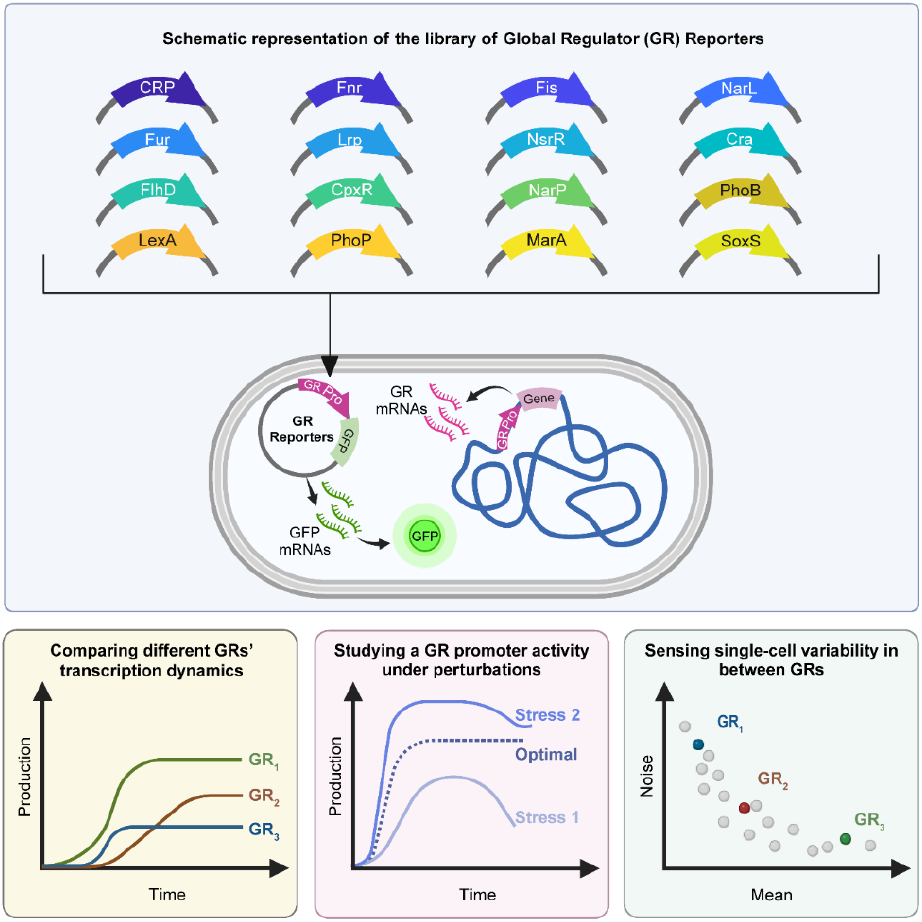

## INTRODUCTION

For millions of years, bacteria have evolved complex transcriptional programs of stress responses. These programs are carried out by transcription factors (TF). While most TFs regulate one to a couple of genes, less than 10% have evolved to directly influence tens to hundreds of genes (Martínez-Antonio & Collado-Vides, 2003). For example, in *E. coli*, 22 TFs alone control 31% of all genes (Supplementary Table S1) (Tierrafría *et al*, 2022). Hereafter, we refer to these 22 TFs as GRs.

In general, GRs participate in complex, life-saving phenotypic changes (Supplementary Table S2) (Gottesman, 1984; Grainger & Busby, 2008). For example, the GR ‘CRP’ regulates hundreds of genes (Tierrafría *et al*, 2022; Zheng *et al*, 2004; Tsai *et al*, 2018) influencing, among other things, how cells use carbon in the absence of glucose (Shimada *et al*, 2011). This, in turn, enhances *E. coli*’s catabolic flexibility during slow, carbon-limited growth (Franchini *et al*, 2015). Similarly, the GR ‘H-NS’ influences approximately two hundred genes (Tierrafría et al., 2022), that particularly affect chromosome compaction (Ueguchi & Mizuno, 1993) during osmotic, pH, and temperature shifts (Arold *et al*, 2010). To study how GRs influence the TFN, one needs to track their levels over time.

Advances in Synthetic Biology (Voigt, 2020; Wang *et al*, 2013; Liu *et al*, 2013) have led to the engineering of two large strain libraries for tracking the expression levels of many genes in live *E. coli* cells. One is the fluorescent transcriptional reporter library (‘TR Library’) (Zaslaver *et al*, 2006). It monitors the transcriptional activity of many natural promoters by having copies of these promoters (in low-copy reporter plasmids) controlling the production of RNA coding for fast-maturing GFP. The other is the ‘YFP fusion library’ (Taniguchi *et al*, 2010). It uses chromosomal integrated YFPs followed by the native genes, both under the control of the natural promoters. Unfortunately, the first library only tracks 7 out of the 22 GRs mentioned above, while the second library only tracks 4 (out of which, 1 is also in the TR Library). Thus, they cannot be used to track most GRs using the same fluorescent protein.

The lack of probes (with the same properties) for all GRs is currently a major limitation in the study of the global transcriptional programs of the TFN of *E. coli* (e.g., for stress adaptation). Specifically, since GRs compose a large fraction of the TFN, tracking their levels over time is necessary to identify the triggers of phenotypic modifications that cells undergo during stress adaptation. For example, in (Baptista *et al*, 2022), tracking σ^38^ levels overtime using a fast-maturing fluorescent protein, allowed establishing that they only change after the beginning of the transition to stationary growth. Tracking more GRs would have allowed determining, e.g., if some of them changed prior to this transition (and, thus, potentially, acted as triggers).

To address this, we present a strain library to measure the single-cell distributions of the transcription dynamics of 16 out of the 22 (arguably) most influential GRs of *E. coli* (Supplementary Table S1). Each strain contains a single-copy plasmid harboring a copy of the native promoter of the GR gene, including the operator sites, to mimic the regulation of the native gene. This promoter controls the transcription of a sequence coding for a fast-maturing GFP variant (Cormack *et al*, 1996). The graphical abstract illustrates the name of the reporters and potential direct applications. Next, we describe how the library was assembled, and show evidence of its functionality in optimal and stressful conditions. Supplementary Figure S1 shows a flowchart of the steps of this study.

## RESULTS

### Assembly and testing of the library of GR Reporters

We first searched in RegulonDB (Tierrafría *et al*, 2022) for the number of genes known to be regulated by each TF in *E. coli*. Next, as mentioned, we defined as GRs the 22 TFs with 40 or more target genes each (in total, they control more than 30% of the genes of *E. coli*).

We also collected the DNA sequences controlling the GRs expression profiles (from RegulonDB). These are the DNA sequences downstream of the previous gene coding region and upstream of the coding region of our GR’s gene of interest. Noteworthy, these regions include the promoters, but do not include codes for any polypeptide. Moreover, no promoters are known to control GR expression outside of these regions. We thus used these DNA sequences as the ‘bioparts’ (Figure 1A_1_) of our GR reporters, which we designed on pBAC plasmids using SnapGene (GSL Biotech) (Figure 1A_2_). As a side note, we failed to synthesize reporters for: IHF, ArcA, NtrC and ModE. In detail, first, we failed to find promoter sequences for ModE and NtrC. Moreover, while we found the promoter sequences of IHF and ArcA, they formed stem loops during synthesis (due to the existence of complementary sequences), which hampered their integration into the plasmid backbones.

**Figure 1:**
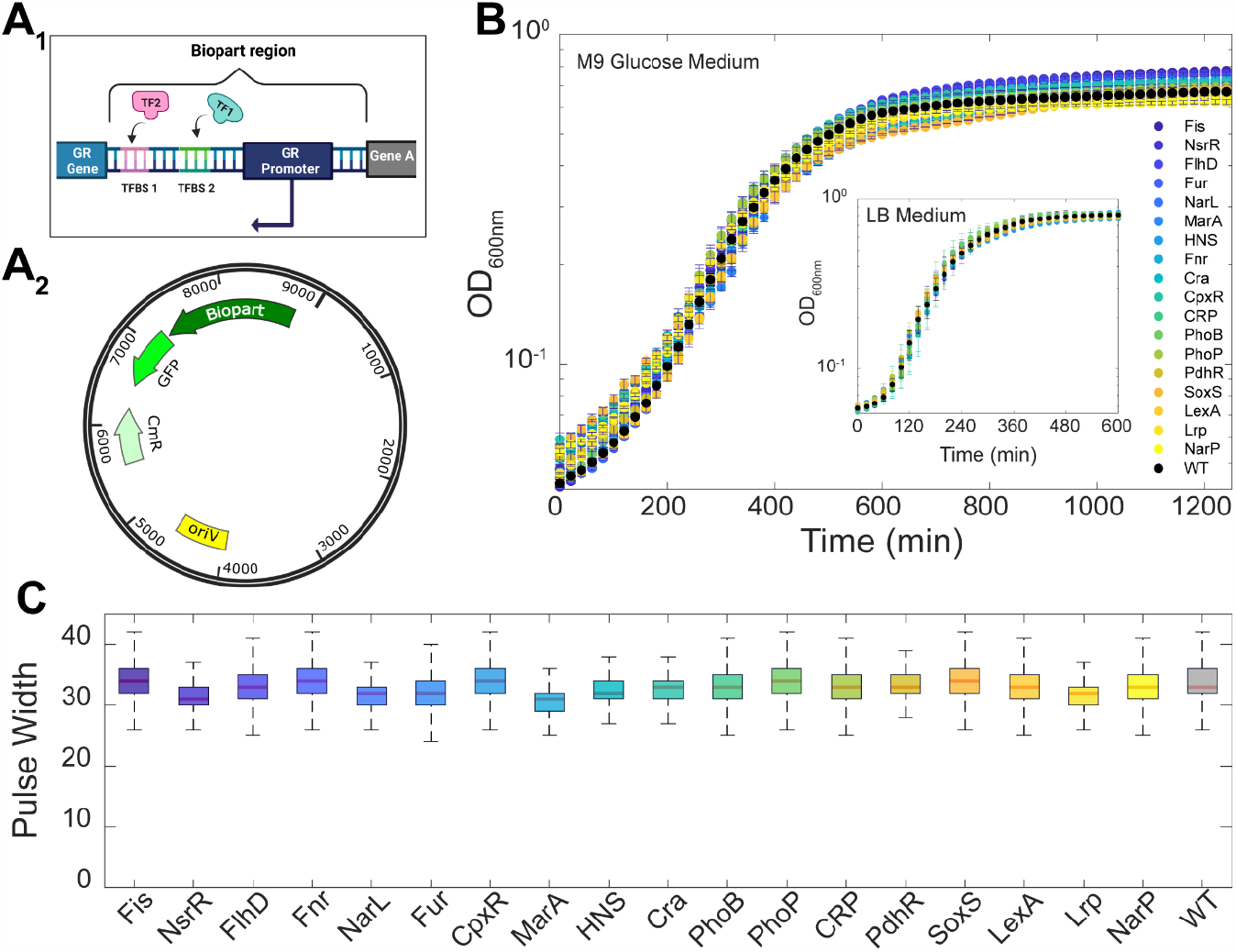
Structure of the plasmids and physiology and morphology of the strains carrying the GR reporters. **(A**_**1**_) Illustration of the region of the DNA from which the biopart was copied from. The biopart is a copy of the coding regions of the natural TF binding sites along with the promoter of the natural GR. Also shown are the start of the GR gene coding region and the end of the gene upstream from the biopart region, to assist identifying the start and end nucleotides of the biopart. **(A**_**2**_) Illustration of the single-copy plasmid (pBAC) carrying the biopart (SnapGene, GSL Biotech). Shown are the regions coding for the biopart, the GFPmut3 downstream from the biopart, a Chloramphenicol resistance gene (CmR), and the origin of replication (OriV). Image created with BioRender.com. **(B)** Growth curves of the WT strain (MG1655) and of the strains carrying the plasmids coding for the GRs, respectively. The OD_600nm_ was measured every 20 min in minimal (M9 Glucose) medium at 37°C. The inset shows the OD_600nm_ in rich (LB) medium. The error bars are the standard error of the mean (SEM) of 3 biological replicates. **(C)** Box plot of the average pulse widths from flow-cytometry, used as a proxy for cell size, of each strain. The center of the boxes indicates the median. The bottom and top edges of the vertical line indicate the 25^th^ and 75^th^ percentiles, respectively.

The plasmids (constructed in Integrated DNA technology, Iowa, U.S.A.,) also include fluorescent protein coding regions downstream of the biopart to report on the transcription rates in single cells (Figure 1A). Specifically, we used single-copy pBAC plasmids containing GFPmut3 (Lee *et al*, 2016) (a kind gift from J.J. Collins). For a schematic representation of the backbone of this plasmid see Data Availability.

GFPmut3 has a strong ribosome binding site to quickly produce many proteins from each RNA (Lee *et al*, 2016) and each protein is 21 times brighter than wild-type GFP (Cormack *et al*, 1996), resulting in very strong signals. Moreover, GFPmut3 has a very fast maturation time (∼4 min, Table 1 in (Balleza *et al*, 2018)) entailing that its signal strength follows RNA numbers with minimum delay. Further, GFPmut3 has weak photostability (Balleza *et al*, 2018), which mimics well the natural RNA degradation times of *E. coli* (∼2-10 min (Bernstein *et al*, 2002)). Finally, GFPmut3 is nontoxic, e.g., it does not form inclusion bodies and it does not react with natural cell components (Cormack *et al*, 1996; Zaslaver *et al*, 2006), implying that it does not interfere with cellular functioning. All these features combined make GFPmut3 an efficient reporter of RNA numbers over time (rather than of the number of proteins that the RNAs code for).

To confirm past reports informing that GFPmut3 is non-toxic, we studied growth in cells with and without the plasmid in minimal and rich growth medium, respectively (Methods section “Bacterial strains and growth conditions”). In both media, we observed only small differences between our strains, including the wild type (WT) (Figure 1B). Moreover, we observed a relatively fast growth rate in LB medium.

We also did not find differences between cell sizes using pulse width (Figure 1C) from flow-cytometry as a proxy (Cunningham, 1990; Traganos, 1984) (Methods section “Flow cytometry”). Additionally, we also obtained cell sizes of 8 strains, including the WT, by microscopy (Methods section “Microscopy and image analysis”), and again did not find significant differences (Supplementary Figure S2A). Finally, we found no differences in cell viability (calculated as CFU/mL, where CFU stands for colony forming units) between the 8 strains, including the WT (Supplementary Figure S2B).

Next, we tested if the promoters of the GR reporters behave similarly to the natural chromosomal-integrated GR promoters. For that, we confronted the average mRNA levels of the natural GR genes (Methods section “RT-PCR”) with the average GFP levels from the GR reporter plasmids (Methods section “Spectrophotometry”). We found a statistically significant linear correlation between them (Figure 2A). However, 2 of the 18 probes (‘HNS’ and ‘PdhR’) can be classified as outliers (Methods section “Fittings and statistical analysis”). Considering the remaining 16 probes alone, we find a high R^2^ for the linear fit (Figure 2A). Moreover, as expected, the linear fit nearly intercepts the origin of the plot aside from a small deviation. Relevantly, that deviation is not significant since a linear fit crossing the origin still results in a fit that has both a high R^2^ and a p-value < 0.05 (Supplementary Figure S3C). We concluded that the remaining 16 synthetic GR reporters are efficient.

**Figure 2:**
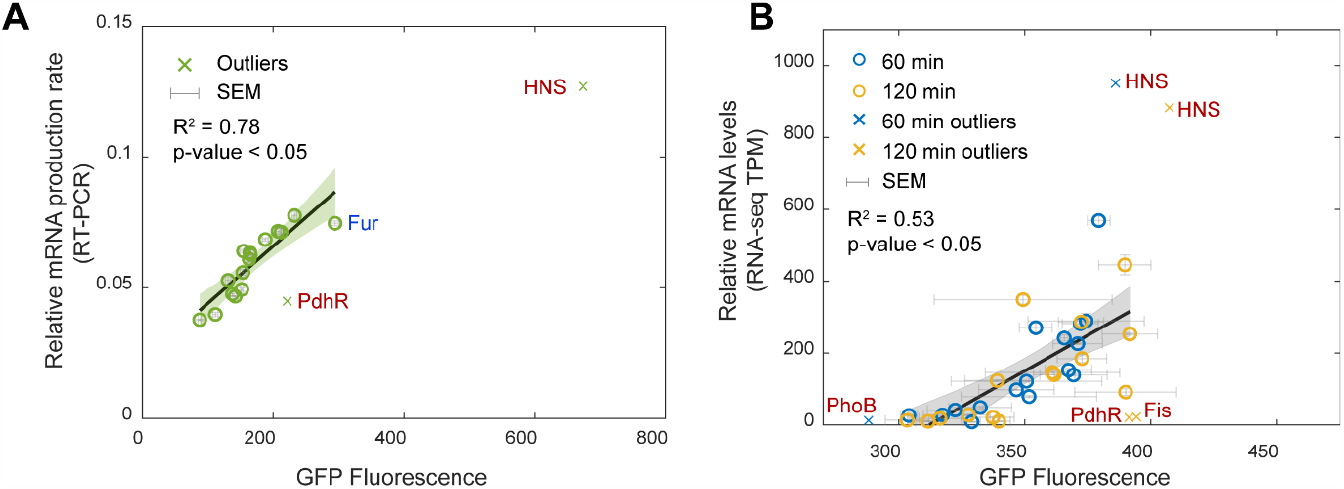
Correlation between RNA levels of the natural genes and the protein expression levels of the plasmids. **(A)** Scatter plot between relative mRNA production rates of native GR chromosomal genes (Methods section “RT-PCR”) and GFP levels of the corresponding reporter plasmids (Methods section “Spectrophotometry”). The error bars are the standard error of the mean (SEM) from 3 biological replicates. Also shown are the p-value and the coefficient of determination, R^2^ (Methods section “Fitting and statistical Analysis”). The names of the outlier datapoints are shown in red, near the crosses. ‘Fur’ is shown in blue (for details see Supplementary Table S3). **(B)** Scatter plot between the TPM normalized counts of GRs from RNA-seq (Methods section “RNA-seq”) and the corresponding GFP. Measurements at 60 and 120 min after placing cells in fresh media (Methods section “Spectrophotometry”). Error bars are the SEM from 3 biological replicates. Also shown are the p-value and the coefficient of determination, R^2^. The x-axis does not start at zero, for easier data visualization. The names of the outlier genes are shown in red, near the crosses.

To further test the efficiency of the GR reporters, we used RNA-seq data from (Almeida *et al*, 2022) for the GRs genes at two time moments during exponential growth (Methods section “RNA-seq”). Then we performed corresponding GRs’ GFP measurements by spectrophotometry in the same growth conditions and time moments. As above, we found a statistically significant linear correlation between the RNA-seq data and the average GFP levels from the GR reporter plasmids (Figure 2B).

Interestingly, again the probes of ‘HNS’ and ‘PdhR’ were outliers (in at least one of the two time points). Due to this confirmation, we did not include these two probes in our library of probes and did not consider them for further analysis. The few other probes classified as outliers were kept in the library since, neither were they outliers when using RT-PCR, nor in the two RNA-seq measurements (60 min and 120 min).

We additionally performed a final statistical analysis of the RT-PCR data (in Figure 2A). Specifically, we considered the effects of removing each of the 16 remaining probes from the data, by assessing if any removal would significantly alter the R^2^ of the linear fit (Figure 2A). The results in Supplementary Table S3 show that only the removal of Fur would increase the R^2^ significantly. However, since Fur was not classified as an outlier in any of the three experiments (RT-PCR and the two RNA-seq time points), we opted for maintaining the strain carrying the probe for Fur in the library. Considering this, along with Figures 2A and 2B, we concluded that the 16 GR reporters are all efficient reporters.

### GR reporter activities during the exponential and stationary growth conditions

We monitored the GRs’ transcription activity in exponential and stationary growth phases (Methods sections “Bacterial strains and stress conditions” and “Spectrophotometry”). To compare the activities between different GR probes, we normalized the raw population fluorescence values (Supplementary Figure S4A) by the corresponding OD_600_ values at the same time points (shown in Figure 1B). This removed the influence from the size of the cell populations. The normalized values were used as a measure of the average single cell fluorescence levels at each time point. We expect these values to differ between strains as they should reflect the expression dynamics of the various GRs.

From Figure 3A, we found a wide variety of behaviors. For example, ‘CRP’ and ‘Fur’ exhibit much higher activity than ‘NarL’ and ‘SoxS’. This is expected since ‘CRP’ and ‘Fur’ are commonly present during exponential growth (Franchini *et al*, 2015; Zhang *et al*, 2005), while ‘NarL’ and ‘SoxS’ are induced during nitrate and oxidative stresses, respectively (Katsir *et al*, 2015; Li & Demple, 1994). Meanwhile, ‘NarL’ and ‘Fis’ decrease, ‘SoxS’ and ‘PhoB’ are stable, and ‘NsrR’ first increases and then decreases, among other diverse behaviors. We expect these differences to arise mostly from differences in RNA production dynamics alone, since all probes share the same RNA degradation rates and the same protein (GFPmut3) translation and degradation rates. The WT strain is not shown as the autofluorescence values are below 30 fluorescence units, which would not be visible (see ‘Data Availability’).

**Figure 3:**
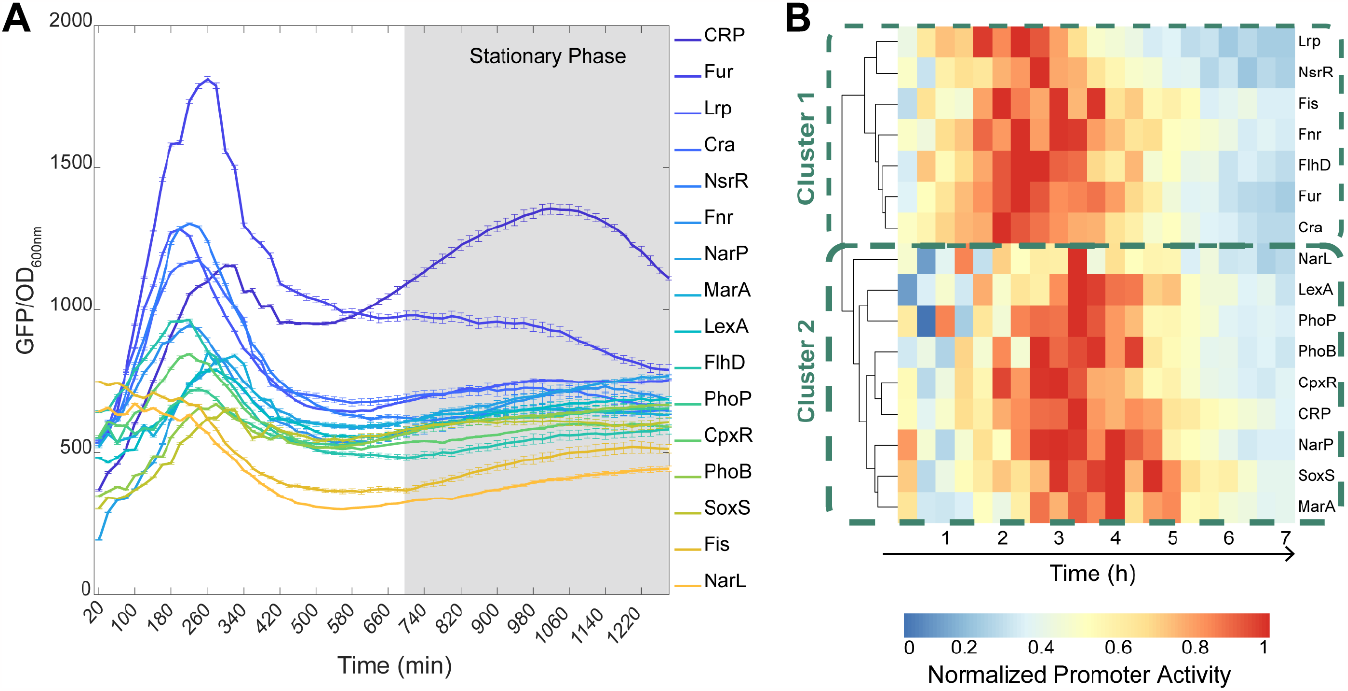
Transcriptional dynamics of the GR reporters in optimal growth conditions. **(A)** Fluorescence levels normalized by the OD_600_ values over time of the strains carrying each of the 16 GR reporter plasmids during exponential and stationary growth phases. Cellular autofluorescence (obtained from the WT strain) was subtracted from all data points, but this had no significant influence on the results (Data Availability). The GR reporters in the legend are listed from top to bottom from the highest to the lowest fluorescence value in the y-axis at the minute 1200, respectively. The grey area is the time window during which cells are in the stationary growth phase, which is preceded by the exponential growth phase. **(B)** Heatmap of the promoter activity 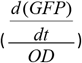 of the reporters of the GRs over time. Obtained using the R package ‘pheatmap’ (version 1.0.12). Values are relative to the maximum level in each GR, to identify when it maximized. High and low values are colored red and blue, respectively (corresponding color bar in the top right corner). The dendrogram is shown on the left side of the heatmap. The GRs were compared using Euclidean distances and were clustered using the complete-linkage method.

We also estimated the changes in GR promoter activities over time (Supplementary Figure S4B and Methods section “Fittings and statistical analysis”). To facilitate comparison, we normalized the intensity levels of the GR reporters over time by their respective maximum expression level. From this, we obtained a heatmap, using the ‘pheatmap’ R-package. Shown in Figure 3B are the first 420 minutes, which is the period of time during which the promoters’ activity changes significantly (Supplementary Figure S4B). The ‘pheatmap’ package estimates the difference in expression levels between two GR reporters from a Euclidean distance 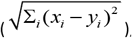, where *x* and *y* are the expression intensity of each GR reporter, respectively. Meanwhile, *i* is the time moment (Supplementary Table S4).

Next, the GR reporters were clustered based on these distances, using a complete-linkage method (pheatmap). The resulting dendrogram in Supplementary Figure S4C shows two dynamical behaviors. The first cluster (*Cluster 1*) has genes whose promoter activity peaks at ∼140 min (Lrp, NsrR, Fis, Fnr, FlhD, Fur and Cra). *Cluster 2* includes genes whose promoter activity peaks 60 min later (NarL, LexA, PhoP, PhoB, CpxR, CRP, NarP, SoxS and MarA).

We also investigated gene ontology (Ashburner *et al*, 2000; Ontology Consortium, 2021) to identify which biological processes are the GR genes of the two clusters involved in. We compared their over-represented biological processes (Supplementary Figure S5, Methods Section “Gene ontology”). Visibly, genes of cluster 1 are associated with positive regulation of biological processes. Meanwhile, genes of cluster 2 are associated to negative regulation of biological processes. Potentially, this could partially explain the differences in promoter activity.

As a side note, the fluorescence levels reported in Figure 3A are affected not only by the transcription activity of the GR reporters, but also by the degradation rates of the RNAs coding for GFP and, by the GFP translation and degradation rates. Nevertheless, these three rates are identical in all strains. Consequently, the fluorescence intensities in Figure 3A should only differ between strains, due to differences in the transcription rate of the GR reporters. Therefore, these intensities are a good proxy for the transcription activities of the GR promoters.

### GR reporters detect differences in single-cell variability of the GRs’ expression level

We measured the variability in the single-cell expression levels of the GR reporters of each strain. In Figure 4A, we plotted the CV^2^ (squared coefficient of variation) against the mean, M, of the single-cell fluorescence levels (Methods section “Flow cytometry”). Then, in agreement with (Taniguchi *et al*, 2010; Bar-Even *et al*, 2006), CV^2^ best fits the data by the ordinary least squares method when assuming the function:

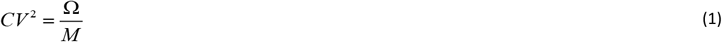

where Ω is a constant (blue curve in Figure 4A).

**Figure 4:**
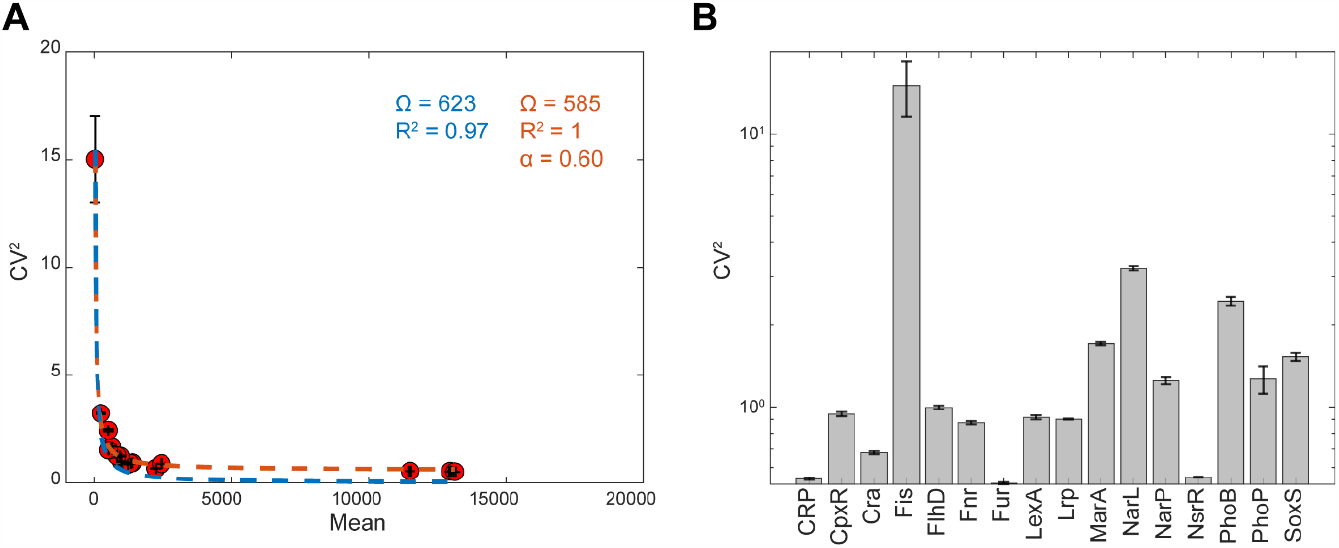
Single-cell expression levels of the GR reporters in optimal growth conditions. **(A)** Scatter plot between the CV^2^ and the mean of the single-cell fluorescence for each strain. Shown are the best fitting lines 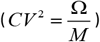 when (red line) and when not (blue line) considering a noise floor (α). Values from cells in the exponential phase (300 min, 0.3 OD_600_ in Figure 3A). **(B)** CV^2^ of single-cell fluorescence levels of the strains carrying each GR reporter in (A). The y-axis is in log scale.

Unfortunately, this model fails to fit well the empirical data for high values of M (M > 2000, Figure 4A). Likely, this is because the model in (1) does not capture the contribution from extrinsic noise sources (Taniguchi *et al*, 2010; Bar-Even *et al*, 2006). These sources include variability in RNAP and ribosome numbers. For example, recent studies reported that single-cell RNAP numbers can differ by ∼45% in cells growing in the same conditions, specifically in the exponential and stationary growth phases (Baptista *et al*, 2022). We thus corrected the model by adding a constant *α* (representing a lower noise limit). This improves the fit, as measured by R^2^ (red curve in Figure 4A):

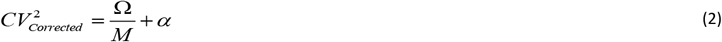

Note that 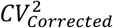 is only an estimation of the *CV*^2^ of the natural RNAs of the GRs. Specifically, we expect these natural RNA numbers of the GRs to be subject to different noise sources than the RNAs of the probes. For example, the natural genes are chromosome-integrated (unlike the reporters). Because of this, we expect positive supercoiling buildup to be influential in natural RNA numbers, but not in the RNA numbers of the probes (Chong *et al*, 2014).

Nevertheless, the reporters contain the entire promoter regions of the natural GRs. As such, they likely have inherited specific features of the dynamics of transcription of the natural promoters, controlling not only the mean rate of production (Figures 2A and 2B), but also the single-cell variability in the RNA production dynamics (Figure 4B).

To test this, we also measured the CV^2^ of single-cell proteins levels of the two natural, chromosomal integrated GRs (Methods sections “Bacterial strains and stress conditions”) that are present in the YFP fusion library (Taniguchi *et al*, 2010). Visibly, the single cell variability in expression levels of the ‘CpxR’ gene is higher than of the ‘Fur’ gene in both the chromosome integrated YFP strains (4× higher, as measured by CV^2^) as well as when it is plasmid borne (2× higher). The difference in the relative increase could be due to different noise sources, e.g., positive supercoiling buildup. As a side note, we did not compare the CV^2^ of the natural genes (YFP strain library) and the reporters (presented here) of each GR, because the fluorescent proteins that they are tagged with differ.

### The GR reporters exhibit specificity and sensitivity to weak stresses

We considered the strains containing ‘Fur’, ‘MarA’, ‘SoxS’, and ‘LexA’ GR reporters, which are responsive to stresses caused by an excess of iron, tetracycline, hydrogen peroxide, and Kanamycin, respectively (Supplementary Table S2) (Methods section “Bacterial strains and stress conditions”).

Specifically, first, ‘Fur’ (ferric uptake regulator) binds to DNA and represses transcription in the presence of iron (Chen *et al*, 2007). This, evidence suggests, contributes to maintaining iron homeostasis (Latorre *et al*, 2018). Adding excess iron to the media should increase the transcriptional activity of the GR reporter of Fur. Meanwhile, ‘MarA’, coded by the ‘marRAB’ operon, controls the expression of genes involved in the resistance to antibiotics, including tetracycline (Sulavik *et al*, 1995). Thus, tetracycline should induce the GR reporter of MarA (it would be possible to use aTc instead, if at lower concentrations). Similarly, oxidative stresses, caused by an excess of hydrogen peroxide (H_2_O_2_), damages *E. coli* cells (Rodríguez-Rojas *et al*, 2020). One gene known to be induced by H_2_O_2_ is ‘SoxS’, which regulates the expression of several genes (Li & Demple, 1994). We should observe an increase in SoxS GR reporter’s activity under oxidative stress. Finally, we expect Kanamycin to increase the activity of the GR reporter of ‘LexA’, since Kanamycin causes DNA damage (Kang *et al*, 2012), which activates the SOS response pathway (D’Ari, 1985).

We measured, using spectrophotometry, the time-lapse response intensities of our reporters for ‘Fur’, ‘MarA’, ‘SoxS’, and ‘LexA’ to all four stresses above, respectively. We first tested weak stress conditions, i.e., not strong enough to affect cell growth rates significantly (Supplementary Figure S7). For this, we started by normalizing the fluorescence values of the genes at each time point by the corresponding OD_600_ values of their strains at the same time point (i.e., GFP/OD). Then, we calculated the fold-change in these normalized values, for each GR probe, in each stress condition. For this, we divided these values by the corresponding values when in optimal growth conditions (reported in Figure 3A).

This allows comparing the transcription rates of the GRs. A normalized fold-change (FC) of a GR during the stress will be either 1 (in case of no change), or more/less than 1 in case of overexpression/inhibition, respectively. We only used data from 50 min onwards after placing cells under stress, to account for protein production times in *E. coli* (Methods Section “Response times of the GR Reporters”). From Figures 5A_1_-5A_4_, each GR reporter responds effectively to its specific stress, and only that one.

**Figure 5:**
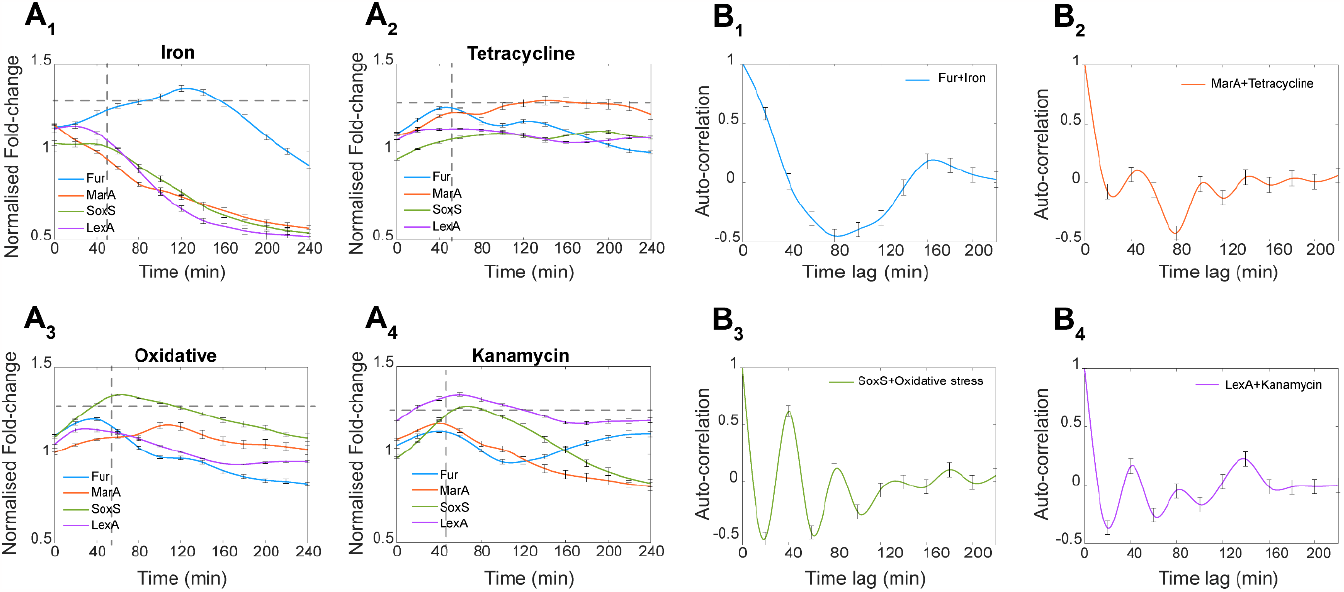
Activities of the GR reporters under weak stress conditions. **(A)** GR reporter levels over time when under weak stresses. Levels are relative to the optimal condition in the same time point. Shown are the stresses: **(A**_**1**_) iron excess (0.2 μM), **(A**_**2**_) Tetracycline (0.05 μg/mL), **(A**_**3**_) oxidative stress (0.6 mM hydrogen peroxide [H_2_O_2_]), and **(A**_**4**_) Kanamycin (25 μg/mL). The error bars are the SEM of 3 biological replicates. Cellular autofluorescence (obtained from the WT strain) was subtracted from each data points, including the optimal condition. Values above 1 correspond to overexpression, while below 1 correspond to repression, when compared to the control. The horizontal dashed lines signal the value above which the fold-change is high enough to be classified as ‘true positive’. Meanwhile, the vertical dashed lines correspond to the time after which we quantify sensitivity and specificity (40 min). **(B)** Autocorrelation of the time-lapse mean expression levels of the GR reporters, when under their respective stress. **(B**_**1**_) iron excess, **(B**_**2**_) Tetracycline, **(B**_**3**_) oxidative stress, and **(B**_**4**_) Kanamycin. The error bars correspond to the SEM of three biological repeats.

From this data, to quantify the reporters’ efficiency, we calculated their sensitivity and specificity (Methods Section “Sensitivity, Specificity, and Fitness”), i.e., their capacity to detect the true signal (sensitivity), while not responding to other signals (specificity), respectively. For this, we define a threshold for the FC, above which the signal is classified as ‘responsive’, while below it is ‘non-responsive’. Specifically, we tested lower and uppear bound threshold values of 1.0 and 1.3, respectively (Supplementary Table S5). A threshold of 1.3 mildly maximizes the fitness (Methods Section “Sensitivity, Specificity, and Fitness”), defined as being the sum of the average sensitivity and average specificity of the 4 reporters to the 4 stresses (Supplementary Figure S6B). Nevertheless, the fitness is not very influenced by the threshold, which suggests that the reporters are functionally robust. Overall, given the high sensitivity and specificity (Supplementary Table S5), we conclude that the reporters are efficient under weak stress conditions.

As a side note, during iron excess, the reporters for ‘MarA’, ‘SoxS’, and ‘LexA’ responded negatively. This is expected, as noted above, and does not affect the sensitivity and specificity. Also noteworthy, the rates of RNA and protein degradation (as well as translation) of the reporters should not differ between strains, with the control, nor in each individual stress condition (although they may differ between stress conditions). Thus, the signals from the GR reporters in each stress can be directly used to compare the transcriptional activities of the GRs in the same stress condition. However, they should not be loosely used to directly compare the transcription activities of the same GR in different stresses. For this, it may be necessary to account for changes in several parameters, including RNA and/or protein degradation rates, etc.

### The GR reporters can detect negative autoregulation mechanisms

Out of the 22 GRs studied, 5 have known positive autoregulation mechanisms, while 8 are known to be subject to negative autoregulation (Tierrafría *et al*, 2022). These mechanisms cause distinct behaviors (e.g., negative autoregulation usually causes oscillations, within certain ranges of parameter values). We tested whether oscillations can be detected by the probes of ‘Fur’, ‘SoxS’, and ‘LexA’, which are negatively autoregulated. As a control, we also evaluated ‘MarA’, which is positively autoregulated (Tierrafría *et al*, 2022). Thus, it should not oscillate.

We searched for oscillations in the time-lapse responses to the weak stress conditions (Figures 5A_1_-5A_4_) (Methods section ‘Fittings and Statistical Analyses’). From (Figures 5B_1_-5B_4_), we find oscillations in ‘Fur’ (80 min. long), ‘SoxS’ (40 min. long), and ‘LexA’ (40 min. long). Meanwhile, we did not find it in ‘MarA’. We conclude that the probes detect negative autoregulation, when existing.

### The GR reporters maintain their specificity and sensitivity under strong stresses

We studied the behavior of the reporters in cells under strong stresses. We classified a stress as ‘strong’ if the growth rate differs from the optimal condition. The protocols to impose the stresses are described in Methods sections “Bacterial strains and stress conditions”. Since these stresses affected the growth rates (Supplementary Figure S8A_1_), we opted for *not* using spectrophotometry to measure the activity of the reporters. Instead, we used flow cytometry to measure reporter activities at the single-cell level. As a side note, in agreement with past reports (Kortman *et al*, 2012; Appenzeller *et al*, 2005), we found faster growth under strong iron excess.

Growth rates have been related with cell size during optimal growth conditions and, thus, could be related with protein levels (Kratz & Banerjee, 2023; Zatulovskiy *et al*, 2022). Thus, we first investigated the average cell size and single-cell fluorescence under the strong stresses. From Supplementary Figure S8A_2_ and S8A_3_, respectively, both are disturbed by the stresses. Thus, we searched for correlations between average cell size, cell fluorescence, and cell growth rate of the strains under the optimal and stress conditions. We found no correlations (Supplementary Figure S8B_1_, S8B_2_, and S8B_3_, see also Supplementary Figure S6A, S6C). We also found no correlations when subjecting each strain to the other stresses (Supplementary Figure S6D). Moreover, at the population level, the differences in mean expression levels of the GR reporters shown in Supplementary Figure S8A_3_ are not, on average, related to changes in mean cell size and/or in mean growth rate. Instead, as expected, they are likely the outcome of the GR reporter’s responsiveness to the stresses.

Next, we evaluated the reporters’ sensitivity at the single cell level, after 120 min under the strong stresses (Methods section “Sensitivity, Specificity, and Fitness”). From (Figures 6A-6D), both in optimal and in strong stress conditions, there is a positive, statistically significant correlation between the reporters’ fluorescence intensity and the pulse width. This is expected since these fluorescence proteins are decaying largely due to cell division (i.e., dilution). However, the fluorescence levels are stronger under stress (specifically the average FITC-H are higher by 1.3 times, 1.5 times, 1.4 times, and 1.3 times, respectively). This causes the inclination of the best-fitting line to be ∼10 times smaller in stress conditions (Figures 6A-6D). As expected, the p-value comparing the slopes (using a t-test statistics) is smaller than 0.05, from which we conclude that the two slopes are not from the same distribution. Overall, we find that all reporters were ‘sensitive’ to the strong stresses, with their response not being affected by small differences in average cell sizes between the strains.

**Figure 6:**
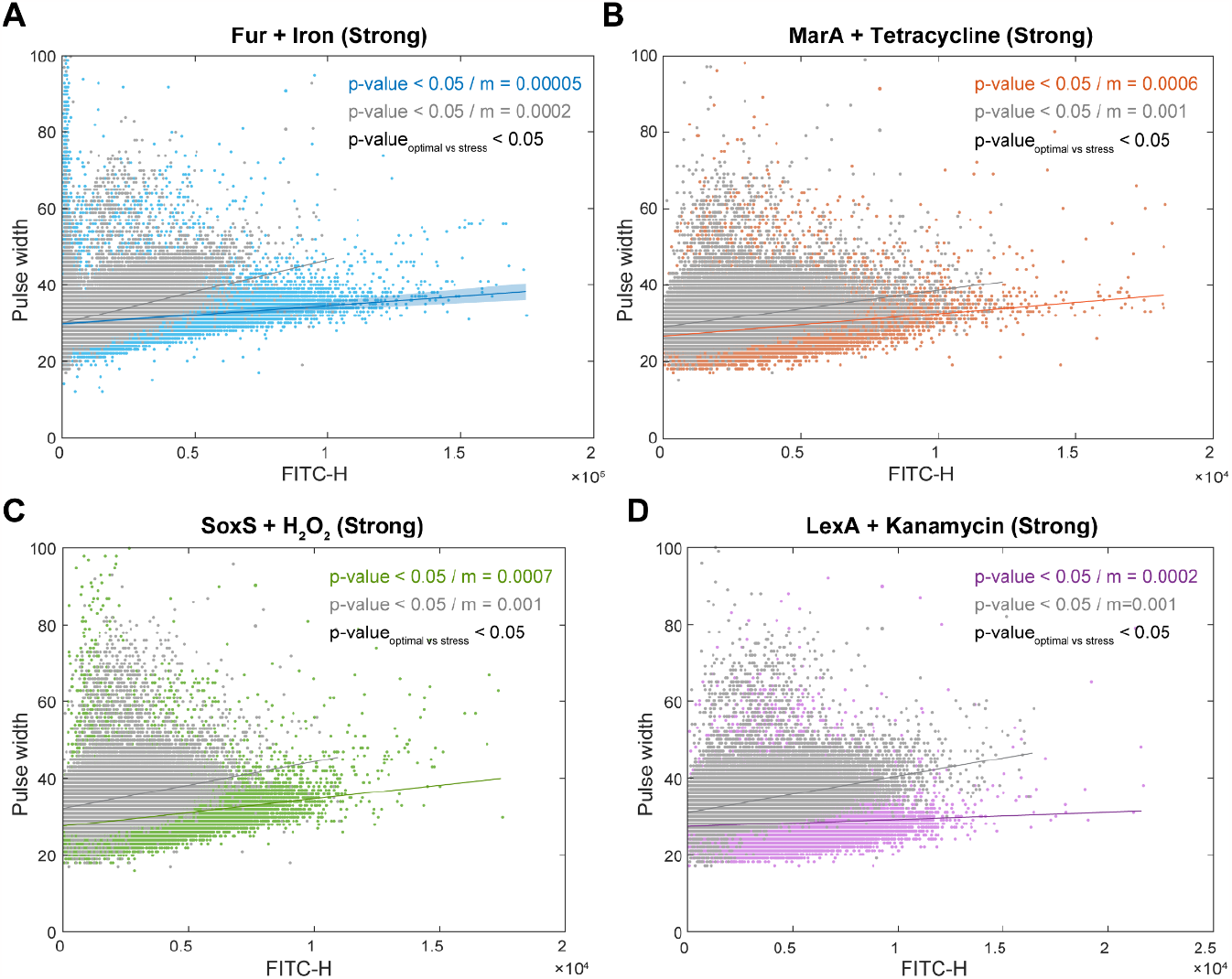
Single-cell size and GR reporter activities under the specific strong stresses. Measurements under **(A)** iron excess (2 μM), **(B)** tetracycline (5 μg/mL), **(C)** oxidative stress (5 mM of hydrogen peroxide), and **(D)** Kanamycin (50 μg/mL). Single-cell expression levels (FITC-H) are plotted against the pulse width (a proxy for cell size). Colored dots are cells subject to the stress. Grey dots are cells in optimal conditions, shown for comparison. Also shown are the best-fitting lines to each of the two cohorts. The *p*-value of each line is from a statistical test of whether the linear fit differs from a horizontal line, using ‘fitlm’ in MATLAB. For p-value > 0.05, we cannot conclude that the line differs from a horizontal line. Measurement time corresponds to 120 min in Supplementary Figure S8A_1_.

Finally, we assessed the reporters’ ‘specificity’ (Methods section “Sensitivity, Specificity, and Fitness”), by subjecting the strains to strong stresses, other than the stress they should detect. Their response strengths were weak in all cases, causing the relationship between pulse width and cell fluorescence to differ little from when in optimal conditions (Supplementary Figure S9). In agreement, in general, the inclinations of the best fitting lines for optimal and for ‘other’ stress conditions are of the same order of magnitude. Also, as expected, average pulse widths and average cells fluorescence of the strains remain uncorrelated, when subject to ‘other’ stresses (Supplementary Figure S6D). We conclude that the GR reporters have high specificity, even under strong stress conditions.

## DISCUSSION

The TFN of *E. coli* has evolved an out-degree distribution that follows a power law (Almeida *et al*, 2022), due to which a small fraction of the genes expressing TFs (22 of 216) directly regulate many genes (∼31% of all ∼4747 genes in the genome (RegulonDB 11.1) (Tierrafría *et al*, 2022). Following the RNA and/or protein levels of each of these GRs is, thus, a key step for dissecting the transcriptional programs of stress response. With this goal, we engineered a library with 16 out of these 22 GR reporters.

We showed that the single-copy plasmids coding for the reporters, under the control of copies of the natural GR promoters, do not affect significantly neither the morphology nor the physiology of the host cells. Additionally, due to the fast maturation times of the GFP coded, the GR reporters track well the native RNA levels of the GRs. As such, the GR reporters can sense changes in the global transcriptional programs a few minutes after these emerge. This is important since, in *E. coli*, most regulatory mechanisms of gene expression act on transcription (Browning & Busby, 2004, 2016; Ruff *et al*, 2015).

Prior to this library, reporters have been engineered for only a few GRs. Also, they differ in their rates of protein folding and degradation, in photo bleaching, etc., hindering direct comparisons (and in assessing how these rates change with environmental conditions, etc.). Our library is the first to allow direct comparisons of the dynamics of most GRs, due to using the same fluorescent reporter and WT strain to engineer all strains.

One expected application of this library is the study of the temporal levels of the GRs during life-threatening, genome-wide stresses affecting hundreds of genes. This may allow explaining how the GRN of *E. coli* carries ordered, short- and long-term transcriptional programs involving large gene cohorts that introduce advantageous phenotypic changes, while robustly maintaining homeostasis.

Moreover, since the reporters are in plasmids, it will be easy to combine them with other synthetic libraries. For example, combining our GR reporters with strains from the KEIO collection (Baba *et al*, 2006) and the ASKA library (Kitagawa *et al*, 2005) should allow assessing how much a given gene influences the GRs. Similarly, a ‘Tunable TF library’(Parisutham *et al*, 2022) was recently produced to synthetically tune the concentration of most TFs. However, it cannot monitor natural production rates. Combining our reporters with this library, one can assess how TFs can affect GR levels, etc.

Interestingly, while the GR reporters have the native promoter sequences (including operator sites), they all exhibited similar response times to the various stresses, as well as to the transition to stationary growth phase. While this is enhanced by all reporters coding for the same GFP protein, it also requires similar transcription rates. As such, our results suggest that there may exist an evolved synchrony in response times between GRs that may be critical for the proper functioning of complex transcriptional programs involving hundreds of genes.

In the future, to enhance this library’s ability to reproduce the temporal numbers of natural GRs, we could introduce their native RBS sequences in front of the GFP coding sequence. We could also use mutant GFP sequences with different production times to account for different protein folding times of the GRs. Moreover, we could use tandem promoters to further fine tune GFP production (Chauhan *et al*, 2022), among other.

Finally, we expect this new library of GR reporters to have wide applicability in synthetic biology, therapeutics, and bioindustrial efforts involving *E. coli*. E.g., the library could be used to investigate if and how GRs are involved in the loss of functionality that synthetic circuits usually suffer with changing external conditions. Second, we can use the library to identify GRs (and subsequent transcriptional programs) involved in bacterial tolerance to antibiotics, which is a critical step for the emergence of resistance. Specifically, we can now track GR numbers, to find which ones may be involved in enhancing tolerance. Finally, this library can be useful in bioindustrial processes. Currently, many bioindustrial processes are energetically costly. E.g., several bioindustrial processes need to be carried out at high temperatures. Potentially, by modifying key transcriptional programs, it may be possible to engineer strains that require lesser extreme temperatures, albeit at the cost of some survivability. Tracking GR numbers is a necessary step for identifying which GRs need to be tuned to adjust those cellular decisions.

## METHODS

### Bacterial strains and growth conditions

We used *E. coli* K-12 MG1655 cells (WT strain), as it is also used in (Zaslaver *et al*, 2006; Taniguchi *et al*, 2010; Parisutham *et al*, 2022). Chemically competent (CC) *E. coli* K-12 MG1655 cells were prepared for plasmid transformation. For each strain, 5 μL of the plasmid DNA (10 ng) coding for the GR reporter was mixed with 50 μL of MG1655 CC (1:10 ratio), and then incubated on ice for 30 min. Next, we kept the mixture at 42 °C on a water bath for 1 minute. Finally, 800 μL of SOC medium was added to the mixture, which was kept at 37 °C under aeration at 250 RPM for 1 hour.

From each mixture, 200 μL was plated using the spread plate method on fresh LB agar plates (2%), prepared by supplementing with antibiotics (34 μg/mL Chloramphenicol). Finally, the plates were kept overnight at 37 °C. The next day, three colonies were picked from each plate and inoculated in fresh LB medium, supplemented with antibiotics (34 μg/mL Chloramphenicol). Afterwards, the cells were incubated at 30 °C overnight with shaking at 250 RPM. The resulting culture cells were diluted into fresh M9 media (0.03 optical density, i.e., OD_600_) and were supplemented with amino acids, vitamin solutions and 0.4% glucose as the carbon source.

In general, measurements were conducted using overnight culture cells. The control condition was M9 medium at 37 °C. On average, the mid-exponential phase was reached in 300 min (OD_600nm_ ∼ 0.3). At this stage, for each of the 18 strains, we performed various measurements of GR reporter activities and other control measurements using RT-PCR, colony counting for assessing cell viability, flow-cytometry, and microscopy.

For weak stress conditions, upon reaching OD_600nm_∼0.3, four strains containing ‘Fur’, ‘MarA’, ‘SoxS’, and ‘LexA’ plasmids, respectively, were subject to iron, Tetracycline, Kanamycin, and H_2_O_2_, respectively. To subject cells to excess of iron, M9 medium was supplemented with iron citrate, as in (Abdul-Tehrani *et al*, 1999). A stock solution of iron citrate (iron-to-citrate ratio, 1:100) was made by dissolving ferrous sulfate (4 mM) in sodium citrate (400 mM). The pH was adjusted to 7 with NaOH. Finally, 0.2 μM of iron citrate was supplemented to the growth medium. Meanwhile, tetracycline and Kanamycin were added to the growth medium in the concentrations of 0.05 μg/mL (Pourahmad Jaktaji & Ebadi, 2013) and 25 μg/mL (Kang *et al*, 2012), respectively. Moreover, oxidative stress was induced by adding 0.6 mM hydrogen peroxide (H_2_O_2_) to the solution. For this, we diluted H_2_O_2_ solution from Merck life sciences (originally at 30%) until it reached 6%, as described in (Imlay & Linn, 1987).

Similarly, for strong stress conditions, the same four strains, upon reaching OD_600nm_∼0.3, were subject to iron (2 μM), Tetracycline (5 μg/mL), Kanamycin (50 μg/mL), and H_2_O_2_ (5 mM H_2_O_2_), respectively. Flow-cytometry measurements were taken 120 min afterwards. Finally, we also diluted overnight culture cells into fresh LB media to monitor the growth as well as to measure their GFP levels after 60 and 120 minutes. The latter was compared to the RNA-seq data from (Almeida *et al*, 2022).

### Spectrophotometry

We measured the optical density at 600 nm (OD_600nm_) to monitor the cell growth, using a Biotek Synergy HTX Multi-Mode Reader. With the same machine, we measured GFP fluorescence of cell populations over time. For this, the solution containing the cells was excited at 485 nm, while the emission was recorded at 528 nm.

### Microscopy and image analysis

To prepare cells for microscopy, first, the cells were pelleted and re-suspended in ∼100 μL of the media. Then, 3 μL of cell suspension were placed on a 2% agarose gel pad made up of M9 medium and kept in between the round microscope slide and a coverslip. It took less than 5 minutes to move cells from the incubator to the microscope and start the observation. This time gap includes the assembly of the microscope imaging chamber containing the slides with the cells. Phase-contrast images were taken by an external phase-contrast system, and then were analyzed by the ‘CellAging’ software (Häkkinen *et al*, 2013) to automatically segment cell borders and extract cell areas. We used the areas as proxies for cell size.

### RT-PCR

Cells were harvested from 3 independent colonies in the mid-exponential phase (0.3 OD_600nm_). Then, 5 mL of the culture was immediately treated with a double volume (10 mL) of RNA protect bacteria reagent (Qiagen, Germany) for 5 minutes at room temperature, to prevent RNA degradation. Next, enzymatic lysis was performed with Tris-EDTA Lysozyme (15 mg/ml) buffer (pH 8.3). From the lysates, the RNA content was isolated using RNeasy purification kit (Qiagen) as per the manufacturer instructions. The RNA yield (∼2 μg/μL) and absorbance ratios A_260_/A_280 nm_ and A_260_/A_230 nm_ were measured by a NanoDrop 2000 Spectrophotometer (Thermo Fisher Scientific, USA). The ratio was found to be 2.0–2.1, which indicates highly purified RNA.

Afterwards, we removed DNA contamination. For this, samples were treated with DNaseI (Thermo Scientific, USA) as per the manufacturer instructions. The cDNA was synthesized from 1 μg of RNA using iScript Reverse Transcription Supermix (Biorad, USA) as per the manufacturer instructions. Next, cDNA samples (10 ng/μl) were mixed with qPCR master mix with iQ SYBR Green supermix (Biorad, USA) with primers (10 μM) for target and reference (16S rRNA) gene, respectively.

Finally, we analyzed the samples using a Biorad MiniOpticon Real Time PCR System (Biorad, USA). The thermal cycling protocol was 40 cycles of 95 °C for 10 s, 52 °C for 30 s, and 72 °C for 30 s, with the fluorescence being read after each cycle. qPCR efficiencies of these reactions were > 95%. No-RT and no-template controls were used to crosscheck for non-specific signals and contamination. Cq values from the CFX ManagerTM Software were used to calculate the fold-change in the target genes (after subtracting for the reference gene).

### Flow cytometry

We measured single-cell fluorescence using an ACEA NovoCyte Flow Cytometer (ACEA Biosciences Inc., San Diego, USA). Cells were diluted (1:10000) into 1 mL of phosphate buffer saline (PBS) solution, vortexed for 10 seconds. For each strain, we obtained 3 biological replicates (50,000 cells each). The flow rate was set to 14 μL/minute. The data was collected by a Novo Express software vs. 1.6 (ACEA Biosciences Inc.).

To detect GFP and YFP, we used a blue laser (488 nm) for excitation. For emission, we set a core diameter of 7.7 μM and a PMT voltage of 600 and we obtained the values of the fluorescein isothiocyanate detection channel (FITC-H) (530/30 nm filter) and of SSC-H (side scattering height) and FSC-H (forward scattering height).

We set two lower bounds (detection thresholds), one for FSC-H (set to 5000) and another for FITC-H (we removed the 1% highest values). These bounds removed any significant interferences from the data. Noteworthy, we never detected significant differences between the 3 repeats in any strain. We also measured the pulse width, which we used as a proxy for cell size (Cunningham, 1990; Traganos, 1984).

### RNA-seq

The RNA-seq data used was first published in (Almeida *et al*, 2022). Briefly, overnight cultured cells were diluted into fresh LB media. RNA-seq measurements were performed 60 and 120 minutes after that. From the raw data in (Almeida *et al*, 2022), RNA sequencing reads were trimmed to remove adapters and low-quality reads by Trimmomatic v.0.36. Quality check was done using fastqc. Trimmed reads were then aligned to the reference genome of *E. coli* MG1655 (NC_000913.3) using Bowtie2 aligner v.2.3.5.1. Unique gene hit counts were calculated with featureCounts from the Rsubread R package (v.1.34.7). Genes with less than 5 counts in more than 3 samples were removed from further analysis. The relative abundance of mRNA in each condition was calculated using Transcripts per Million Normalization (Li & Dewey, 2011).

### Fittings and statistical analysis

To find best fitting lines (e.g., Figures 2A and 2B), we used linear regression functions (MATLAB). Goodness of fit was estimated from R^2^ values. Meanwhile, we used the *t*-statistic with the null hypothesis that the slope of the best fitting line (of a scatter plot between the variables) is not different from zero. For *p*-values smaller than 0.05, we rejected the null hypothesis. Also, we used an iterative procedure to discard outliers in linear correlations (Zaslaver *et al*, 2006) (Figures 2A and 2B). The outliers were classified as data points beyond the boundaries of ellipse that encompassed 90% of the data (Supplementary Figure S3A, S3B).

For calculating a GR promoter activity (as described in (Zaslaver *et al*, 2006)), OD and GFP curves were time shifted, so that all OD curves reached 5% of their maximal OD at the same time after background subtraction. Each background GFP (WT) value was then subtracted from the reporter strain GFP value at the same OD (and not necessarily at the same time point). Expression profiles were calculated by dividing GFP by OD. Then, promoter activity was calculated by taking the time derivative of GFP/OD between consecutive time points 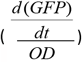. All analysis steps were performed automatically using MATLAB software. The standard error of the mean (SEM) for FC analysis was calculated using an error propagation method, using the formula: 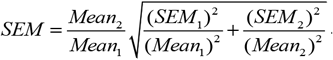. Here, 1 and 2 represent the two groups compared (e.g., stress and control, respectively).

Finally, we searched for the existence of oscillations (an expected outcome of negative gene expression autoregulation) in spectrophotometry data. For this, we applied the ‘detrend’ function (MATLAB) to the average cell population fluorescence levels normalized by population size (OD). ‘detrend’ removes (here, by subtracting) the effect of a best straight-fit line from the data caused by, e.g., the increase in total cell fluorescence due to increasing cell population size. Then, we apply the ‘xcov’ function to the treated data, to calculate the autocorrelation as a function of the lag, at each time point.

### Gene Ontology (GO)

To study the GO representation (Ashburner *et al*, 2000; Ontology Consortium, 2021) of different clusters of GR genes, we performed an overrepresentation test using the PANTHER Classification System (Mi *et al*, 2019). This test finds statistically significant overrepresentations using Fisher’s exact test, which rejects the null hypothesis that there are no associations between the genes’ cohort and the corresponding GO of the biological process for p-values < 0.05. This p-value is corrected for the False Discovery Rate (FDR) using the Benjamini-Hochberg procedure (Benjamini & Hochberg, 1995). In case of biological processes that were overrepresented in both clusters, we opted to classify them in the cluster that has a greater fold enrichment i.e., the cluster that has the most GR genes associated to a biological process than expected by random chance.

### Response times of the GR Reporters

The GR reporters (Supplementary Table S2) operate by producing fluorescent protein GFP at a certain rate once the promoter that they carry is activated by the respective signal. This production is preceded by several events, which take a significant amount of time.

First, once the substance to report (e.g., antibiotic) is introduced in the media, it quickly becomes homogeneously distributed (because the media is under constant mixing). Then, these molecules need to reach the cell cytoplasm and the promoter of interest within. For this, they need to cross the periplasm, and interact with the promoter controlling the expression of the GR reporter. Past studies suggest that substances such as IPTG and Arabinose can take, on average, 20-40 mins to be in sufficient numbers inside the cytoplasm to activate/repress the target promoters efficiently (Megerle *et al*, 2008; Tran *et al*, 2015). We assume an average time length of 30 minutes, for simplicity, for all substances tested here.

Next, once the promoter of the reporter is active, we expect transcription elongation of the RNA coding GFP to be a relatively fast process. Specifically, the DNA sequence coding for GFPmut3 is 717 nucleotides long (Lee *et al*, 2016). Meanwhile, transcription elongation should occur at approximately ∼42 nuc /s (Phroskin *et al*, 2010). Thus, elongation should take ∼15-20 seconds. Finally, it follows translation and maturation of the reporter protein, GFPmut3, which is ∼4 minutes (Balleza *et al*, 2018).

Given the above, we estimate that our GR reporter, if its promoter is sensitive to the substance, will have already increased sufficiently in numbers approximately ∼50 minutes, after the target substance is added to the media.

### Sensitivity, Specificity, and Fitness

To define the sensitivity, specificity, and fitness of the GR reporters, first, we considered the data from spectrophotometry (Figures 5A-5D). For weak stresses (which do not disturb growth rates or cell sizes), GFP and OD measurements were conducted using spectrophotometry. We define that if the fold-change between two conditions in the expression level of a GR reporter is above a specific threshold, then the signal is ‘positive’. Else, if the fold-change is equal or below the threshold, then the signal is negative.

Moreover, we classify signals as “true” or “false”. For this, we use Supplementary Table S2. For e.g., the GR ‘Fur’. should respond to iron excess. Also, Fur should not respond to the other stresses tested (excess of H_2_O_2_, Tetracycline, and Kanamycin), since there is no past evidence of such responsiveness. Given this, if the reporter for Fur responds positively to iron excess, we will classify its signal as ‘true positive’. Else, we classify its signal as ‘false negative’. Meanwhile, if it responds positively to another stress, we classify that signal as false positive. Else, we classify that signal as true negative.

Given this, the ‘sensitivity’ of a GR reporter is defined as its capacity to detect true signals. Formally, it can be quantified by a ‘true positive rate’ (TPR), which is the number of true positive (TP) findings over the number of all positive (P) findings (true positive plus false positive findings):

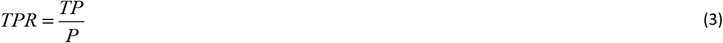

When having time series data on a reporter from a spectrophotometer, we apply this formula to each time moment, and then obtain the average over all time moments, t, as follows:

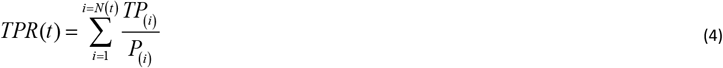

Meanwhile, ‘specificity’ of a reporter is its capacity to remain unresponsive when there is no true signal to detect. In general, specificity is quantified by a ‘true negative rate’, which is the fraction between true negatives (TN) and total negatives (N), i.e., true, and false negatives:

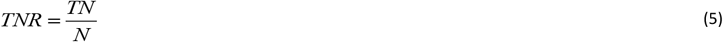

Having information on several time points from a spectrophotometer, one can use the formula:

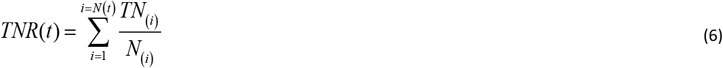

Finally, we define fitness as the average between the sensitivity and specificity. Assuming several time points, it follows:

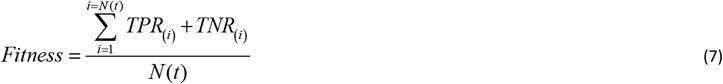

For flow-cytometry data, there is only one time point. Moreover, the definition of TP, TN, FP, and FN differ. In this case, we compare the inclinations of the best fitting lines to the scatter plots between FITC-H (a proxy for transcription rates) and pulse width (a proxy for cell size).

Assume an inclination *m*_*o*_ for a population of cells carrying the plasmid responsive to stress *x*, when in the optimal condition. Now, assume an inclination *m*_*x*_ for cells under the ‘correct’ stress condition. Notably, if the reporter is to be responsive to a stress, FITC-H value should increase. Meanwhile, pulse width should not differ or differ little. This will cause m_x_ to be smaller than m_o_.

Thus, i) if *m*_*x*_ < *m*_*o*_ for the cells carrying reporter X under the appropriate stress X, then it is a TP; ii) if *m*_*x*_ > *m*_*o*_ for the cells carrying reporter X under a stress other than X, then it is a TN; iii) if *m*_*x*_ < *m*_*o*_ for the cells carrying reporter X under a stress other than X, then it is a FP; and, iv) if *m*_*x*_ > *m*_*o*_ for the cells carrying reporter X under a stress X, then it is a FN.

## ACKNOWLEDGEMENTS

We thank Jane and Aatos Erkko Foundation [10-10524-38 to A.S.R.]; Sigrid Jusélius Foundation [230181 to A.S.R.]; Suomalainen Tiedeakatemia (to S.D. and R.J.); EDUFI Fellowship [TM-21-11655 to R.J.]; Finnish Cultural Foundation [00222452 and 00212591 to I.S.C.B.]; Tampere University Graduate Program (to V.C.). The funders had no role in study design, data collection and analysis, decision to publish, or preparation of the manuscript. We thank Antti Häkkinen for his helpful feedback on the statistical analysis. Finally, the authors acknowledge the support of the Biocenter Finland, Tampere Flow Cytometry Facility, and Tampere Imaging Facility.

## DATA AVAILIBILITY

Experimental datasets will be made available on Dryad upon Manuscript acceptance.

## AUTHOR CONTRIBUTIONS

A.S.R. and S.D. conceived the study. A.S.R. supervised the study. S.D. designed the constructs, planned, and executed all measurements. R.J. planned and executed most data analysis. V.C. and S.M.D.O. contributed to the design of the constructs. V.K. contributed to the RT-PCR measurements. S.D. and I.S.C.B. contributed to some data analysis. A.S.R. and S.D. drafted all documents, which were revised by all co-authors.

## SUPPLEMENTARY DATA

Supplemental Information includes 9 supplementary figures, and 5 supplementary Tables and can be found with this article online.

## LIBRARY DISTRIBUTION

The library of GR reporters will be available for distribution in Addgene.

## INCLUSION AND DIVERSITY

We support inclusive, diverse, and equitable conduct of research.

## REFERENCES

Abdul-Tehrani H, Hudson AJ, Chang YS, Timms AR, Hawkins C, Williams JM, Harrison PM, Guest JR & Andrews SC (1999) Ferritin mutants of Escherichia coli are iron deficient and growth impaired, and fur mutants are iron deficient. J Bacteriol 181: 1415–1428

Almeida BLB M Bahrudeen MN, Chauhan V, Dash S, Kandavalli V, Häkkinen A, Lloyd-Price J S D Cristina P, Baptista ISC, Gupta A, et al (2022) The transcription factor network of E. coli steers global responses to shifts in RNAP concentration. Nucleic Acids Res

Appenzeller BMR, Yañez C, Jorand F & Block J-C (2005) Advantage provided by iron for Escherichia coli growth and cultivability in drinking water. Appl Environ Microbiol 71: 5621–5623

Arold ST, Leonard PG, Parkinson GN & Ladbury JE (2010) H-NS forms a superhelical protein scaffold for DNA condensation. Proc Natl Acad Sci U S A 107: 15728–15732

Ashburner M, Ball CA, Blake JA, Botstein D, Butler H, Cherry JM, Davis AP, Dolinski K, Dwight SS, Eppig JT, et al (2000) Gene Ontology: tool for the unification of biology. Nat Genet 25: 25–29

Baba T, Ara T, Hasegawa M, Takai Y, Okumura Y, Baba M, Datsenko KA, Tomita M, Wanner BL & Mori H (2006) Construction of Escherichia coli K-12 in-frame, single-gene knockout mutants: the Keio collection. Mol Syst Biol 2: 2006.0008-2006.0008

Balleza E, Kim JM & Cluzel P (2018) Systematic characterization of maturation time of fluorescent proteins in living cells. Nat Methods 15: 47–51

Baptista ISC, Kandavalli V, Chauhan V, Bahrudeen MNM, Almeida BLB, Palma CSD, Dash S & Ribeiro AS (2022) Sequence-dependent model of genes with dual σ factor preference. Biochim Biophys Acta Gene Regul Mech 1865: 194812

Bar-Even A, Paulsson J, Maheshri N, Carmi M, O’Shea E, Pilpel Y & Barkai N (2006) Noise in protein expression scales with natural protein abundance. Nat Genet 38: 636–643

Benjamini Y & Hochberg Y (1995) Controlling the False Discovery Rate: A Practical and Powerful Approach to Multiple Testing. J R Stat Soc Series B Stat Methodol 57: 289–300

Bernstein JA, Khodursky AB, Lin P-H, Lin-Chao S & Cohen SN (2002) Global analysis of mRNA decay and abundance in Escherichia coli at single-gene resolution using two-color fluorescent DNA microarrays. Proceedings of the National Academy of Sciences 99: 9697 LP – 9702

Browning DF & Busby SJW (2004) The regulation of bacterial transcription initiation. Nat Rev Microbiol 2: 57–65

Browning DF & Busby SJW (2016) Local and global regulation of transcription initiation in bacteria. Nat Rev Microbiol 14: 638–650

Chauhan V, Bahrudeen MNM, Palma CSD, Baptista ISC, Almeida BLB, Dash S, Kandavalli V & Ribeiro AS (2022) Analytical kinetic model of native tandem promoters in E. coli. PLoS Comput Biol 18: e1009824

Chen Z, Lewis KA, Shultzaberger RK, Lyakhov IG, Zheng M, Doan B, Storz G & Schneider TD (2007) Discovery of Fur binding site clusters in Escherichia coli by information theory models. Nucleic Acids Res 35: 6762–6777

Chong S, Chen C, Ge H & Xie XS (2014) Mechanism of transcriptional bursting in bacteria. Cell 158: 314–326

Cormack BP, Valdivia RH & Falkow S (1996) FACS-optimized mutants of the green fluorescent protein (GFP). Gene 173: 33–38

Cunningham A (1990) Fluorescence pulse shape as a morphological indicator in the analysis of colonial microalgae by flow cytometry. J Microbiol Methods 11: 27–36

D’Ari R (1985) The SOS system. Biochimie 67: 343–347

Franchini AG, Ihssen J & Egli T (2015) Effect of global regulators RpoS and cyclic-AMP/CRP on the catabolome and transcriptome of Escherichia coli K12 during carbon- and energy-limited growth. PLoS One 10: e0133793

Gottesman S (1984) Bacterial regulation: global regulatory networks. Annu Rev Genet 18: 415–441

Grainger DC & Busby SJW (2008) Global regulators of transcription in Escherichia coli: mechanisms of action and methods for study. Adv Appl Microbiol 65: 93–113

Häkkinen A, Muthukrishnan A-B, Mora A, Fonseca JM & Ribeiro AS (2013) CellAging: a tool to study segregation and partitioning in division in cell lineages of Escherichia coli. Bioinformatics 29: 1708–1709

Imlay JA & Linn S (1987) Mutagenesis and stress responses induced in Escherichia coli by hydrogen peroxide. J Bacteriol 169: 2967–2976

Kang TM, Yuan J, Nguyen A, Becket E, Yang H & Miller JH (2012) The aminoglycoside antibiotic kanamycin damages DNA bases in Escherichia coli: caffeine potentiates the DNA-damaging effects of kanamycin while suppressing cell killing by ciprofloxacin in Escherichia coli and Bacillus anthracis. Antimicrob Agents Chemother 56: 3216–3223

Katsir G, Jarvis M, Phillips M, Ma Z & Gunsalus RP (2015) The Escherichia coli NarL receiver domain regulates transcription through promoter specific functions. BMC Microbiol 15: 174

Kitagawa M, Ara T, Arifuzzaman M, Ioka-Nakamichi T, Inamoto E, Toyonaga H & Mori H (2005) Complete set of ORF clones of Escherichia coli ASKA library (a complete set of E. coli K-12 ORF archive): unique resources for biological research. DNA Res 12: 291–299

Kortman GAM, Boleij A, Swinkels DW & Tjalsma H (2012) Iron availability increases the pathogenic potential of salmonella typhimurium and other Enteric pathogens at the intestinal epithelial interface. PLoS One 7: e29968

Kratz JC & Banerjee S (2023) Dynamic proteome trade-offs regulate bacterial cell size and growth in fluctuating nutrient environments. Commun Biol 6: 486

Latorre M, Quenti D, Travisany D, Singh KV, Murray BE, Maass A & Cambiazo V (2018) The role of Fur in the transcriptional and iron homeostatic response of Enterococcus faecalis. Front Microbiol 9: 1580

Lee JW, Gyorgy A, Cameron DE, Pyenson N, Choi KR, Way JC, Silver PA, Del Vecchio D & Collins JJ (2016) Creating Single-Copy Genetic Circuits. Mol Cell 63: 329–336

Li B & Dewey CN (2011) RSEM: accurate transcript quantification from RNA-Seq data with or without a reference genome. BMC Bioinformatics 12: 323

Li Z & Demple B (1994) SoxS, an activator of superoxide stress genes in Escherichia coli. Purification and interaction with DNA. J Biol Chem 269: 18371–18377

Liu D, Hoynes-O’Connor A & Zhang F (2013) Bridging the gap between systems biology and synthetic biology. Front Microbiol 4: 211

Martínez-Antonio A & Collado-Vides J (2003) Identifying global regulators in transcriptional regulatory networks in bacteria. Curr Opin Microbiol 6: 482–489

Megerle JA, Fritz G, Gerland U, Jung K & Rädler JO (2008) Timing and dynamics of single cell gene expression in the arabinose utilization system. Biophys J 95: 2103–2115

Mi H, Muruganujan A, Ebert D, Huang X & Thomas PD (2019) PANTHER version 14: more genomes, a new PANTHER GO-slim and improvements in enrichment analysis tools. Nucleic Acids Res 47: D419–D426

Ontology Consortium G (2021) The Gene Ontology resource: enriching a GOld mine. Nucleic Acids Res 49: D325–D334

Parisutham V, Chhabra S, Ali MZ & Brewster RC (2022) Tunable transcription factor library for robust quantification of regulatory properties in Escherichia coli. Mol Syst Biol 18: e10843

Phroskin S, Rahmouni R, Mironov A & Nudler A (2010) Cooperation between translating ribosomes and RNA polymerase in transcription elongation. Science 328: 504–508

Pourahmad Jaktaji R & Ebadi R (2013) Study the expression of marA gene in ciprofloxacin and tetracycline resistant mutants of esherichia coli. Iran J Pharm Res 12: 923–928

Rodríguez-Rojas A, Kim JJ, Johnston PR, Makarova O, Eravci M, Weise C, Hengge R & Rolff J (2020) Non-lethal exposure to H2O2 boosts bacterial survival and evolvability against oxidative stress. PLoS Genet 16: e1008649

Ruff EF, Record MT Jr & Artsimovitch I (2015) Initial events in bacterial transcription initiation. Biomolecules 5: 1035–1062

Shimada T, Fujita N, Yamamoto K & Ishihama A (2011) Novel roles of cAMP receptor protein (CRP) in regulation of transport and metabolism of carbon sources. PLoS One 6: e20081

Sulavik MC, Gambino LF & Miller PF (1995) The MarR repressor of the multiple antibiotic resistance (mar) operon in Escherichia coli: prototypic member of a family of bacterial regulatory proteins involved in sensing phenolic compounds. Mol Med 1: 436–446

Taniguchi Y, Choi PJ, Li G-W, Chen H, Babu M, Hearn J, Emili A & Xie XS (2010) Quantifying E. coli Proteome and Transcriptome with Single-Molecule Sensitivity in Single Cells. Science 329: 533–538

Tierrafría VH, Rioualen C, Salgado H, Lara P, Gama-Castro S, Lally P, Gómez-Romero L, Peña-Loredo P, López-Almazo AG, Alarcón-Carranza G, et al (2022) RegulonDB 11.0: Comprehensive high-throughput datasets on transcriptional regulation in Escherichia coli K-12. Microb Genom 8

Traganos F (1984) Flow cytometry: principles and applications. I. Cancer Invest 2: 149–163

Tran H, Oliveira SMD, Goncalves N & Ribeiro AS (2015) Kinetics of the cellular intake of a gene expression inducer at high concentrations. Mol Biosyst 11: 2579–2587

Tsai M-J, Wang J-R, Yang C-D, Kao K-C, Huang W-L, Huang H-Y, Tseng C-P, Huang H-D & Ho S-Y (2018) PredCRP: predicting and analysing the regulatory roles of CRP from its binding sites in Escherichia coli. Sci Rep 8

Ueguchi C & Mizuno T (1993) The Escherichia coli nucleoid protein H-NS functions directly as a transcriptional repressor. EMBO J 12: 1039–1046

Voigt CA (2020) Synthetic biology 2020-2030: six commercially-available products that are changing our world. Nat Commun 11: 6379

Wang Y-H, Wei KY & Smolke CD (2013) Synthetic biology: advancing the design of diverse genetic systems. Annu Rev Chem Biomol Eng 4: 69–102

Zaslaver A, Bren A, Ronen M, Itzkovitz S, Kikoin I, Shavit S, Liebermeister W, Surette MG & Alon U (2006) A comprehensive library of fluorescent transcriptional reporters for Escherichia coli. Nat Methods 3: 623–628

Zatulovskiy E, Lanz MC, Zhang S, McCarthy F, Elias JE & Skotheim JM (2022) Delineation of proteome changes driven by cell size and growth rate. Front Cell Dev Biol 10: 980721

Zhang Z, Gosset G, Barabote R, Gonzalez CS, Cuevas WA & Saier MH Jr (2005) Functional interactions between the carbon and iron utilization regulators, Crp and Fur, in Escherichia coli. J Bacteriol 187: 980–990

Zheng D, Constantinidou C, Hobman JL & Minchin SD (2004) Identification of the CRP regulon using in vitro and in vivo transcriptional profiling. Nucleic Acids Res 32: 5874–5893

