## Supplementary Data for "A library of reporters of the global regulators of gene expression of *Escherichia coli*"

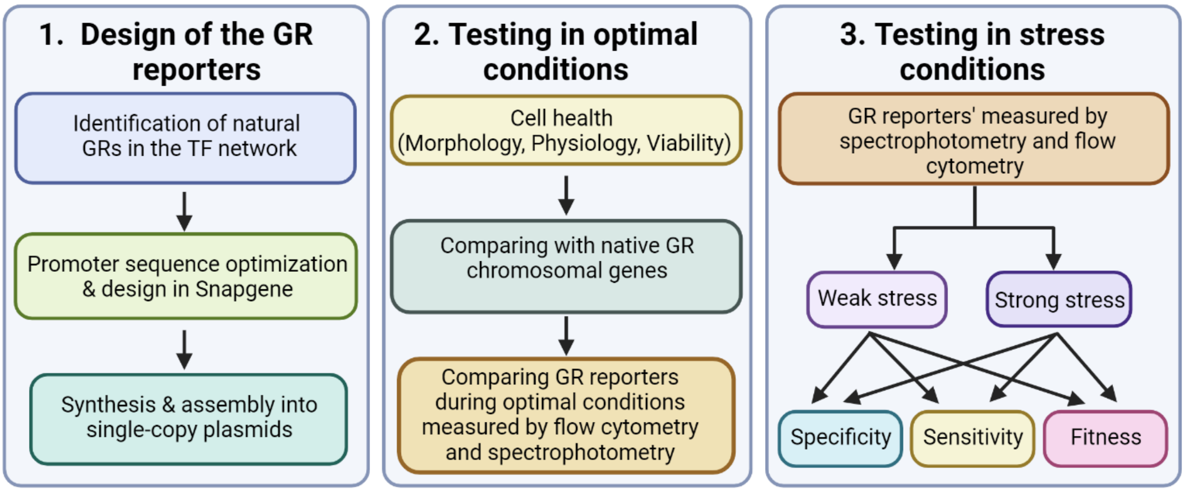


**Figure S1:** Graphical representation of the workflow.

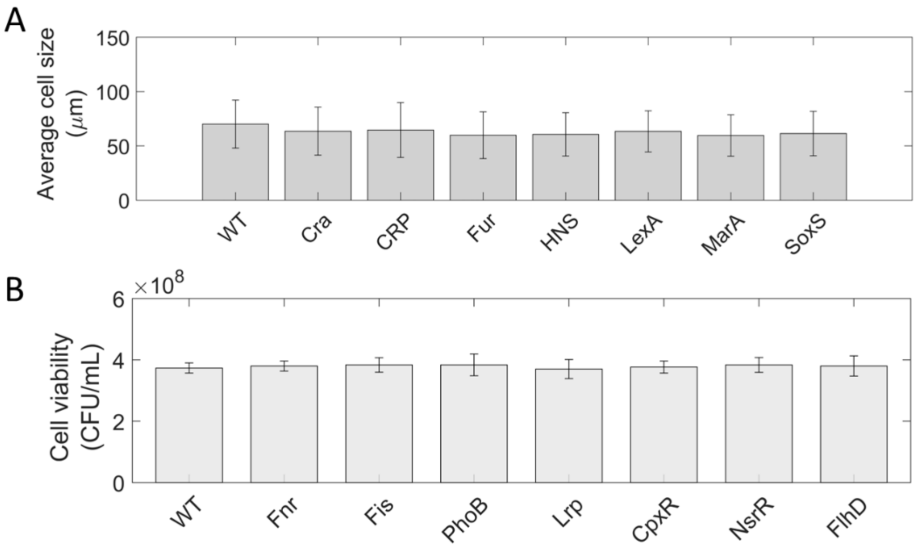


**Figure S2: (A)** Average cell size of several of the strains carrying GR plasmids and the WT (MG1655) strain, as measured by microscopy and image analysis. The error bars are the standard deviation (STDEV). **(B)** Viable cell count (as measured by CFU/mL) of the same strains. Error bars are the standard error of the mean (SEM) of 3 biological replicates.

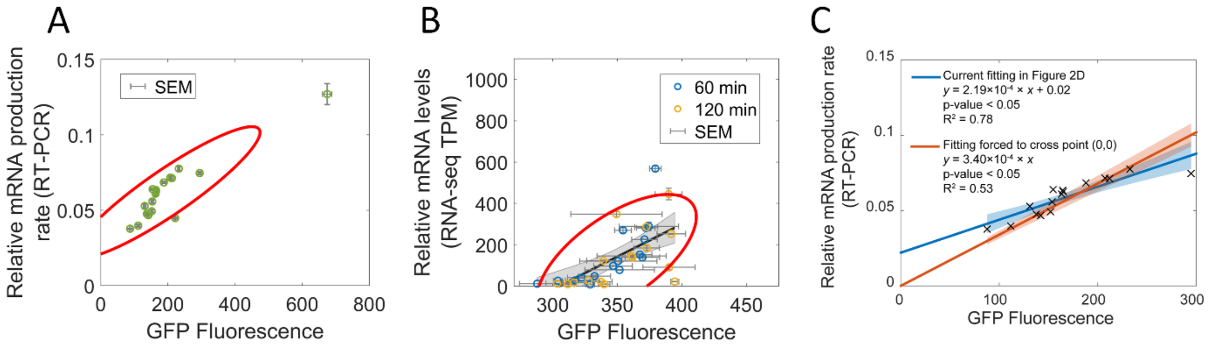


**Figure S3: (A)** Relative mRNA production of the GR reports (as measured by 1/ΔCq values from RT-PCR) plotted against GFP fluorescence measured by spectrophotometry. **(B)** RNA-seq TPM values at 60 min and at 120 min after placing the cells in fresh LB medium plotted against GFP fluorescence measured by spectrophotometry. The red ellipse is the 90% confidence ellipse, which defines the region with 90% of all datapoints. Datapoints in

(A) and (B) that fall outside of the ellipse were classified as being outliers and, therefore, were not considered when establishing the linear fit in Figures 2A and 2B in the main manuscript, respectively. **(C)** Scatter plot between relative mRNA production rates of the 16 native GR chromosomal genes and the GFP levels of the corresponding reporter plasmids (listed in Supplementary Table S1). We added two linear fits. The blue line is the standard linear fit (also shown in Figure 2A in the main manuscript). Meanwhile, the orange line was obtained by forcing the fitting to cross the origin of the plot. For both lines, we show their equation, along with the p-value and coefficient of determination, $R^2$ (Methods section "Fitting and statistical Analysis"). Moreover, the shadow areas of each line represent the 95% confidence intervals of the regression coefficients. These areas overlap for GFP levels ranging from 120 to 300, implying that the lines, in this region, are not statistically distinct.

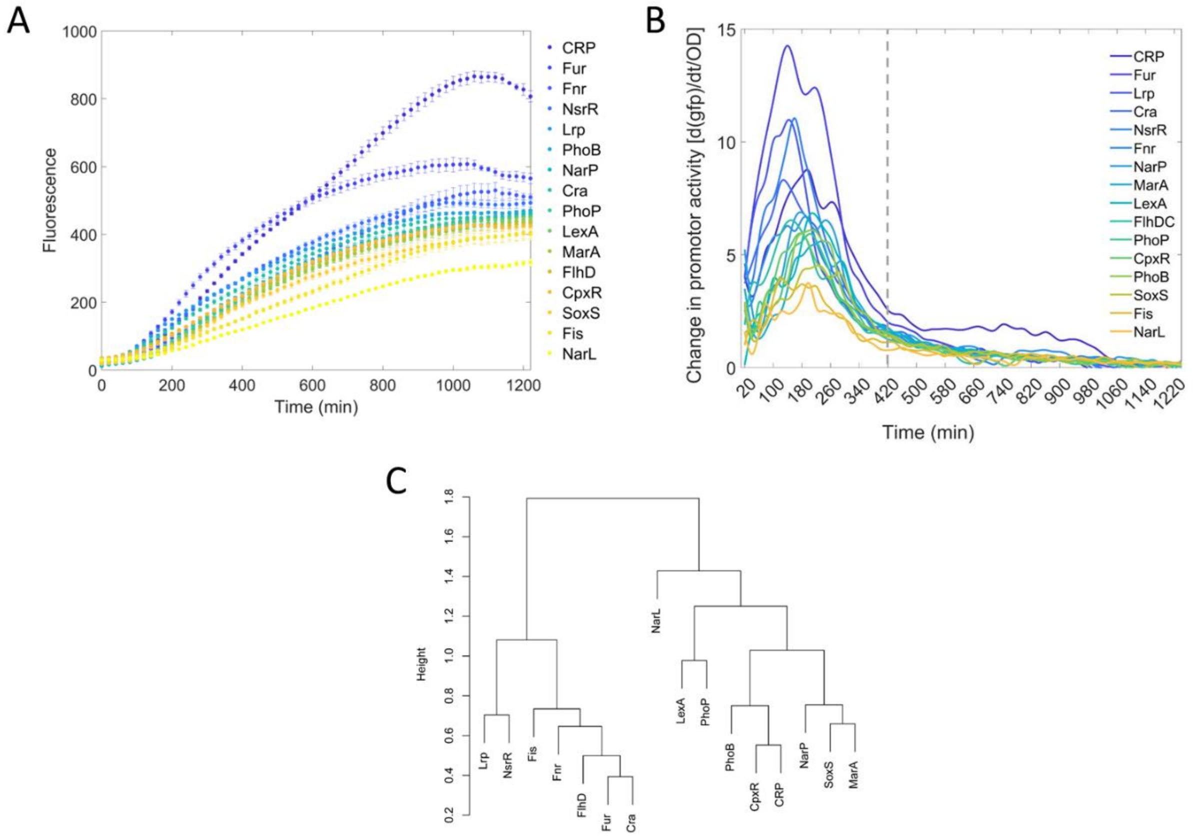


**Figure S4: (A)** Time lapse fluorescence of each of the 16 strains carrying the GR reporters, respectively. Data from when placing the overnight culture in the new medium until the cells reach the stationary growth phase. The average fluorescence of the WT cells over time (named 'cellular background fluorescence' in "Data Availability") was subtracted from the data, but this had no significant influence on the results. **(B)** Change in GR promoter activity ( $\frac{\frac{d(GFP)}{dt}}{OD}$ ) during optimal growth conditions calculated from the fluorescence normalized by $OD_{600}$ in Figure 3A. The dotted vertical line at 420 min marks the moment until when the data was used to obtain the heatmap in Figure 3B of the main manuscript. **(C)** Cluster dendrogram of the GR promoter activity over time, using the 'pheatmap' software R package, based on the empirical data until 420 min.

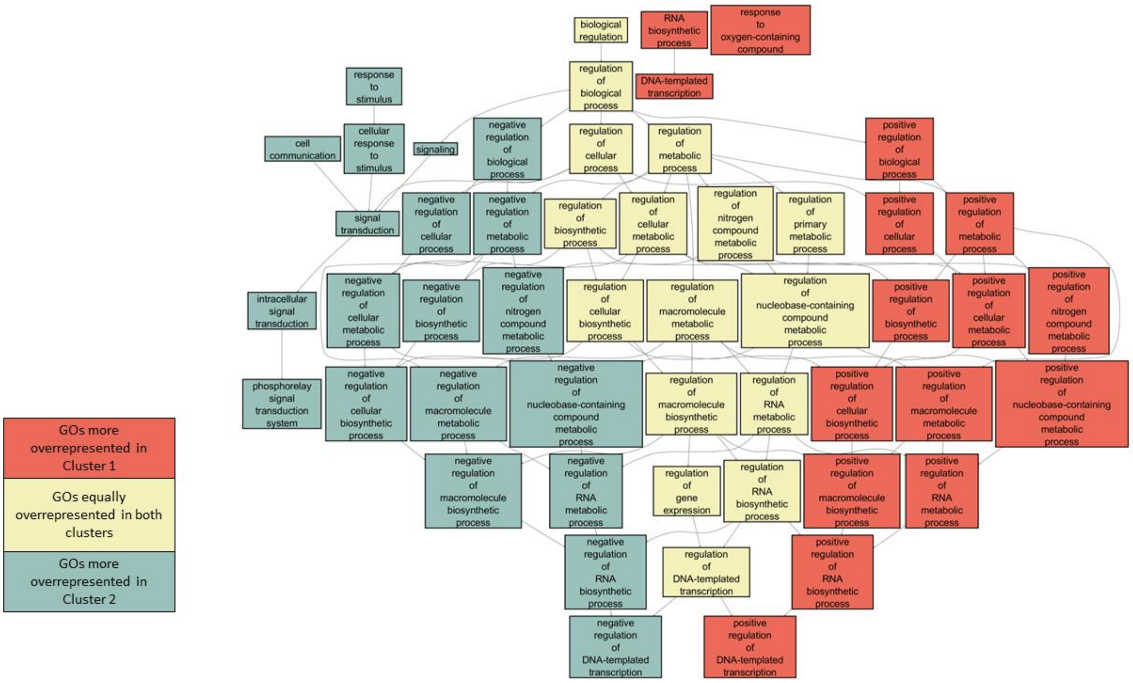


**Figure S5:** Overrepresented gene ontology (GO) terms and their ancestors for the genes controlled by the different GRs. The more general biological processes are connected to specific ontology terms by an arrow pointing to the latter. The ontologies that have the ancestors controlled by cluster 1 and/or 2 are colored differently.

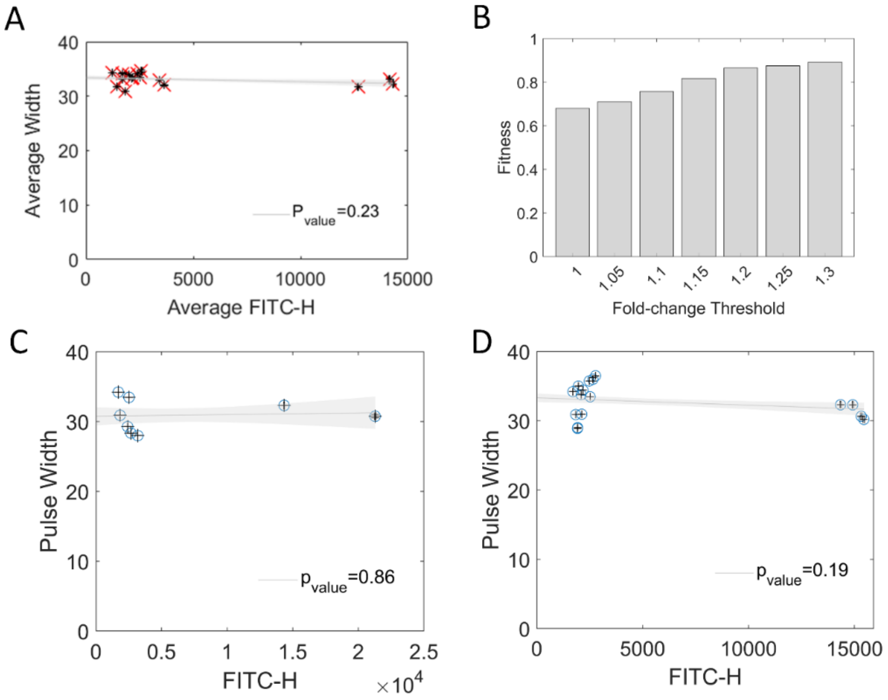


**Figure S6: (A)** Scatter plot of the average single-cell fluorescence (measured by FITC-H) of each strain carrying a GR reporter (data from 16 strains), plotted against the average pulse width of the same cell population (50,000 cells per population) in optimal growth conditions. Also shown are the best fitting line and the p-value from a statistical test of whether the line differs from a horizontal line, using 'fitlm' in MATLAB. For a p-value > 0.05, we cannot conclude that the line differs from the horizontal line. **(B)** Fitness of the GR probes. Fitness is defined as the average between the specificity and the sensitivity values, as a function of a minimum threshold for fold changes, above which the probe is classified as being responsive to the stress. **(C-D)** Scatter plots of the average cell pulse width plotted against the average cell fluorescence (measured by FITC-H) of each strain (Fur, MarA, LexA, and SosX) when **(C)** a given strain is subjected to the specific strong stress that it should respond to (Iron, Tetracycline, Kanamycin, and Oxidative stress, respectively), and, **(D)** when a given strain is subjected to the other three strong stresses that it should not respond to. Each data point is an average from 50,000 cells. Also shown are the best fitting lines and the p-value from a statistical test of whether the linear fit differs from a horizontal line, using 'fitlm' in MATLAB. For a p-value > 0.05, we cannot conclude that the line differs from the horizontal line.

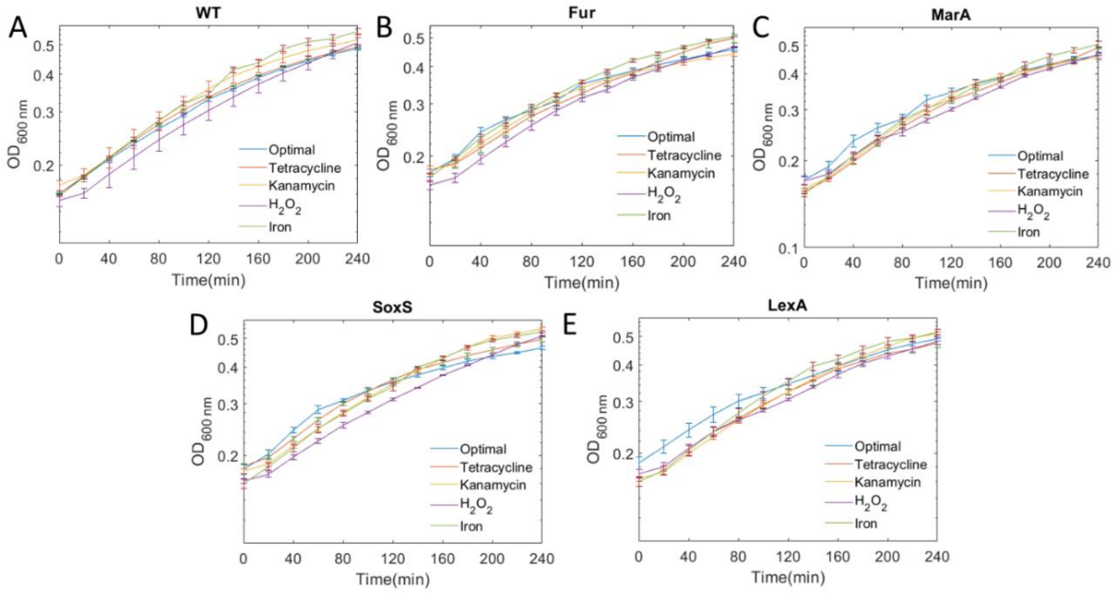


**Figure S7:** Growth curves of the **(A)** WT strain MG1655 and of cells containing GR reporters of **(B)** Fur, **(C)** MarA, **(D)** SoxS, and **(E)** LexA when subject to an external stress once reaching $OD_{600}$ of 0.3 (from "0 min" onwards).

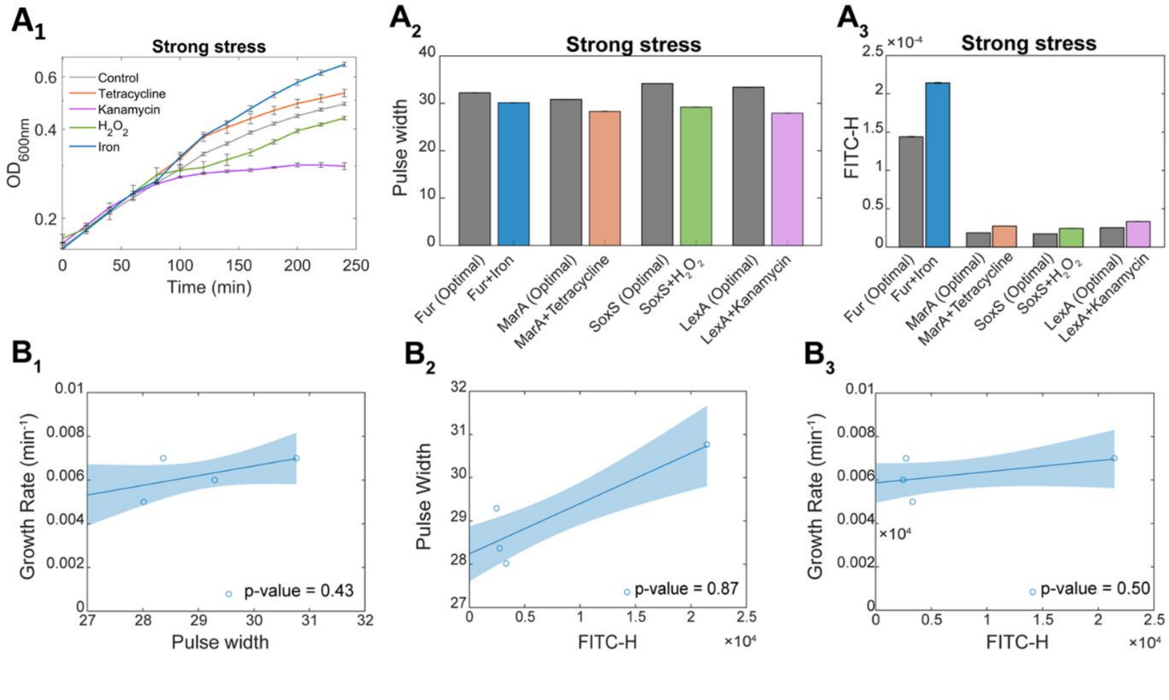


**Figure S8:** **(A₁)** Cell growth rates under strong stresses (Methods Section "Bacterial strains and stress conditions"). **(A₂)** Average pulse width (proxy for cell size). **(A₃)** Average FITC-H (single-cell fluorescence) of the strains carrying the GR reporters expectedly responsive to the stresses. Also shown are the scatter plots of **(B₁)**

average growth rate versus average pulse width, (**B₂**) average FITC-H vs. average pulse width, and (**B₃**) average growth rate vs. average FITC-H of the strains carrying the GR reporters responsive to the respective strong stresses.

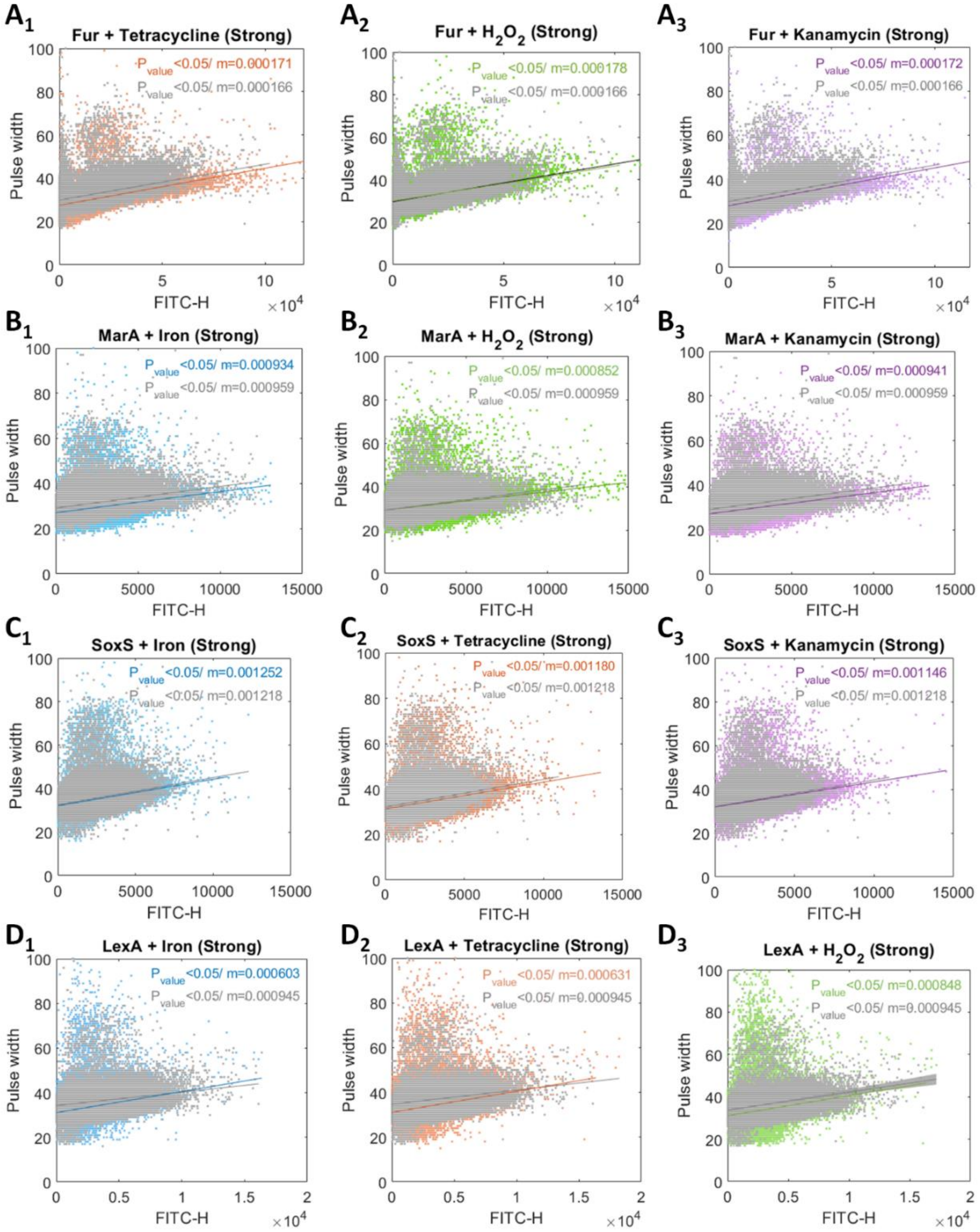


**Figure S9:** Single-cell GR reporter levels under strong stresses. Shown are single-cell expression levels (FITC-H) plotted against the corresponding single-cell pulse widths, measured by flow cytometry. Each figure shows data from cells subject to the stress (colored dots) and from cells not subject to the stress (grey dots). Also shown are the best linear fits to both cell cohorts. The p-values and slopes of the best fitting lines of each cohort are obtained from statistical tests determining whether the linear fits differ from horizontal lines, using the 'fitlm' function in MATLAB. For a p-value > 0.05, we cannot conclude that the line differs from a horizontal line. To compare whether the two slopes of the model are significantly different, a t-test statistic was calculated by

$\frac{m_1 - m_2}{\sqrt{SE_1^2 + SE_2^2}}$, where $m_1$ and $m_2$ are the slopes of the fits to each cohort, respectively, while $SE_1$ and $SE_2$ are the corresponding standard errors, respectively. Finally, to assess if the result is not expected by random chance, a two-tailed *p*-value was calculated. Since the *p*-value was smaller than 0.05 in all cases, we concluded that the two slopes are not from the same distribution.

**Table S1:** List of known global regulators of gene expression in *Escherichia coli* (genes coding for TFs that control 40 or more genes). Also shown are the number of genes that each GR controls (information from RegulonDB as of 19.04.2023) (Tierrafría *et al*, 2022). It follows information on which libraries of synthetic constructs are these GR present in and whether the GR is subject to autoregulation.

| Serial Number | GR names | No of genes directly regulated | No of genes indirectly regulated | GR reporter Library | TR Library (Zaslaver *et al*, 2006) | YFP Fusion Library (Taniguchi *et al*, 2010) | Autoregulation in GRs (Tierrafría *et al*, 2022) |
|---|---|---|---|---|---|---|---|
| 1 | CRP | 606 | 2546 | Yes | yes | | +,- |
| 2 | FNR | 319 | 1139 | Yes | | | - |
| 3 | IHF | 261 | 695 | | | | None |
| 4 | Fis | 245 | 984 | Yes | | | - |
| 5 | HNS | 201 | 441 | | | | None |
| 6 | ArcA | 189 | 437 | | | | None |
| 7 | NarL | 132 | 39 | Yes | | | None |
| 8 | Fur | 138 | 578 | Yes | yes | Yes | - |
| 9 | Lrp | 110 | 141 | Yes | | | - |
| 10 | NsrR | 86 | 169 | Yes | | Yes | None |
| 11 | Cra | 82 | 748 | Yes | | | None |
| 12 | FlhD | 80 | 21 | Yes | | Yes | None |
| 13 | CpxR | 71 | 162 | Yes | | Yes | + |
| 14 | NarP | 66 | 0 | Yes | | | None |
| 15 | PhoB | 65 | 242 | Yes | yes | | + |
| 16 | LexA | 61 | 66 | Yes | yes | | - |
| 17 | PhoP | 59 | 237 | Yes | yes | | +,- |
| 18 | NtrC | 56 | 30 | | | | None |
| 19 | MarA | 52 | 89 | Yes | | | + |
| 20 | ModE | 46 | 132 | | | | None |
| 21 | SoxS | 45 | 270 | Yes | yes | | - |
| 22 | PdhR | 41 | 58 | | yes | | - |

**Table S2:** Biological processes that the GRs are known to affect.

| Serial Number | GR | Biological Function |
|---|---|---|
| 1 | CRP | Glucose starvation (Grainger *et al*, 2005) |
| 2 | FNR | Transition from aerobic to anaerobic metabolism (Unden & Schirawski, 1997) |
| 3 | Fis | DNA-binding protein involved in nucleoid organization (Kelly *et al*, 2004) |
| 4 | HNS | Chromosome compaction (Ueguchi & Mizuno, 1993) |
| 5 | NarL | Anaerobic respiration (Cai *et al*, 2022) |
| 6 | Fur | Global transcription repression as a response to iron excess (Chen *et al*, 2007) |
| 7 | Lrp | Adaptation to changes in the nutritional environment (Landgraf *et al*, 1996) |

| | | |
|---|---|---|
| 8 | NsrR | Nitric oxide-sensitive repressor of transcription (Partridge *et al*, 2009) |
| 9 | Cra | Carbon metabolism in nutrient limited environments (Shimizu, 2013) |
| 10 | FlhD | Flagellum biosynthesis (Gauger *et al*, 2007) |
| 11 | CpxR | Biofilm formation (Dudin *et al*, 2014) |
| 12 | NarP | Anaerobic respiration using nitrate (Brosse *et al*, 2022) |
| 13 | PhoB | Inorganic phosphate metabolism regulation (Marzan & Shimizu, 2011) |
| 14 | LexA | SOS response pathway regulation (D'Ari, 1985) |
| 15 | PhoP | Low $Mg^{2+}$ concentration (Groisman, 2001) |
| 16 | MarA | Antibiotic resistance (Sulavik *et al*, 1995) |
| 17 | PdhR | Pyruvate catabolism regulation (Anzai *et al*, 2020) |
| 18 | SoxS | Superoxide stress protection (Li & Demple, 1994) |

**Table S3:** (Related to Figure 2A in the main manuscript). The table shows the outcome on the linear fitting of removing one data point (gene). Shown are the p-values of *t*-tests of the linear fits (Methods section 'Fitting and Statistical Analysis'). For *p*-values smaller than 0.05, we rejected the null hypothesis. Also shown are the $R^2$'s of the goodness of fit of the line of the remaining 15 data points, after removing the datapoint.

| Gene excluded from the linear fitting | *p*-value | $R^2$ |
|---|---|---|
| CRP | < 0.05 | 0.75 |
| CpxR | < 0.05 | 0.79 |
| Cra | < 0.05 | 0.79 |
| Fis | < 0.05 | 0.76 |
| FlhD | < 0.05 | 0.79 |
| Fnr | < 0.05 | 0.81 |
| Fur | < 0.05 | 0.89 |
| LexA | < 0.05 | 0.78 |
| Lrp | < 0.05 | 0.77 |
| MarA | < 0.05 | 0.79 |
| NarL | < 0.05 | 0.73 |
| NarP | < 0.05 | 0.79 |
| NsrR | < 0.05 | 0.77 |
| PhoB | < 0.05 | 0.79 |
| PhoP | < 0.05 | 0.78 |
| SoxS | < 0.05 | 0.78 |
| None | < 0.05 | 0.78 |

**Table S4:** Euclidean distances between the expression level over time of all pairs of GRs. The distances are scaled by colors from red to white, corresponding to high to low.

| | CpxR | SoxS | Fur | Lrp | NsrR | Cra | FlhD | CRP | Fnr | Fis | NarL | NarP | PhoB | LexA | PhoP | MarA |
|---|---|---|---|---|---|---|---|---|---|---|---|---|---|---|---|---|
| CpxR | 0 | 0.95 | 0.69 | 1.15 | 0.70 | 0.68 | 0.73 | 0.55 | 0.66 | 0.78 | 1.01 | 0.90 | 0.70 | 1.10 | 1.00 | 1.01 |
| SoxS | | 0 | 1.24 | 1.79 | 1.39 | 1.30 | 1.39 | 0.65 | 1.03 | 1.15 | 1.38 | 0.75 | 1.03 | 0.99 | 1.10 | 0.66 |

| | | | | | | | | | | | | | | | | |
|---|---|---|---|---|---|---|---|---|---|---|---|---|---|---|---|---|
| Fur | | | 0 | 0.83 | 0.70 | 0.39 | 0.50 | 0.81 | 0.48 | 0.66 | 1.29 | 1.11 | 0.99 | 1.11 | 1.12 | 1.29 |
| Lrp | | | | 0 | 0.70 | 0.65 | 0.74 | 1.35 | 0.95 | 1.08 | 1.43 | 1.60 | 1.54 | 1.58 | 1.47 | 1.71 |
| NsrR | | | | | 0 | 0.59 | 0.78 | 0.98 | 0.81 | 0.97 | 0.97 | 1.22 | 1.07 | 1.28 | 1.06 | 1.26 |
| Cra | | | | | | 0 | 0.48 | 0.87 | 0.56 | 0.67 | 1.21 | 1.23 | 1.09 | 1.26 | 1.19 | 1.34 |
| FlhD | | | | | | | 0 | 0.86 | 0.65 | 0.62 | 1.34 | 1.17 | 1.03 | 1.20 | 1.28 | 1.37 |
| CRP | | | | | | | | 0 | 0.62 | 0.84 | 1.11 | 0.71 | 0.75 | 0.89 | 0.95 | 0.81 |
| Fnr | | | | | | | | | 0 | 0.73 | 1.26 | 0.97 | 1.08 | 1.03 | 0.97 | 1.17 |
| Fis | | | | | | | | | | 0 | 1.44 | 1.15 | 0.93 | 0.96 | 1.22 | 1.24 |
| NarL | | | | | | | | | | | 0 | 1.43 | 1.14 | 1.33 | 1.37 | 1.21 |
| NarP | | | | | | | | | | | | 0 | 0.97 | 1.06 | 1.03 | 0.71 |
| PhoB | | | | | | | | | | | | | 0 | 0.96 | 1.25 | 1.01 |
| LexA | | | | | | | | | | | | | | 0 | 0.98 | 0.87 |
| PhoP | | | | | | | | | | | | | | | 0 | 0.96 |
| MarA | | | | | | | | | | | | | | | | 0 |

**Table S5:** Sensitivity and specificity of the GR reporters to weak stresses for different threshold values for LFCs.

| Threshold LFC = 1 | | | | | | |
|---|---|---|---|---|---|---|
| **Stress** | **Sensitivity (TPR)** | **Specificity (TNR)** | **TP** | **FP** | **TN** | **FN** |
| Tetracycline | 0.26 | 1.00 | 13 | 37 | 2 | 0 |
| Kanamycin | 0.35 | 1.00 | 13 | 24 | 15 | 0 |
| Oxidative | 0.34 | 1.00 | 13 | 25 | 14 | 0 |
| Iron | 0.52 | 0.97 | 12 | 11 | 28 | 1 |

| Threshold LFC = 1.05 | | | | | | |
|---|---|---|---|---|---|---|
| **Stress** | **Sensitivity (TPR)** | **Specificity (TNR)** | **TP** | **FP** | **TN** | **FN** |
| Tetracycline | 0.28 | 1.00 | 13 | 33 | 6 | 0 |
| Kanamycin | 0.41 | 1.00 | 13 | 19 | 20 | 0 |
| Oxidative | 0.41 | 1.00 | 13 | 19 | 20 | 0 |
| Iron | 0.65 | 0.94 | 11 | 6 | 33 | 2 |

| Threshold LFC = 1.10 | | | | | | |
|---|---|---|---|---|---|---|
| **Stress** | **Sensitivity (TPR)** | **Specificity (TNR)** | **TP** | **FP** | **TN** | **FN** |
| Tetracycline | 0.43 | 1.00 | 13 | 17 | 22 | 0 |

| | | | | | | |
|---|---|---|---|---|---|---|
| Kanamycin | 0.50 | 1.00 | 13 | 13 | 26 | 0 |
| Oxidative | 0.52 | 0.97 | 12 | 11 | 28 | 1 |
| Iron | 0.71 | 0.92 | 10 | 4 | 35 | 3 |

| Threshold LFC = 1.15 | | | | | | |
|---|---|---|---|---|---|---|
| **Stress** | **Sensitivity (TPR)** | **Specificity (TNR)** | **TP** | **FP** | **TN** | **FN** |
| Tetracycline | 0.65 | 0.94 | 11 | 6 | 33 | 2 |
| Kanamycin | 0.68 | 1.00 | 13 | 6 | 33 | 0 |
| Oxidative | 0.67 | 0.92 | 10 | 5 | 34 | 3 |
| Iron | 0.80 | 0.88 | 8 | 2 | 37 | 5 |

| Threshold LFC = 1.2 | | | | | | |
|---|---|---|---|---|---|---|
| **Stress** | **Sensitivity (TPR)** | **Specificity (TNR)** | **TP** | **FP** | **TN** | **FN** |
| Tetracycline | 0.83 | 0.93 | 10 | 2 | 37 | 3 |
| Kanamycin | 0.70 | 0.86 | 7 | 3 | 36 | 6 |
| Oxidative | 0.88 | 0.86 | 7 | 1 | 38 | 6 |
| Iron | 1.00 | 0.87 | 7 | 0 | 39 | 6 |

| Threshold LFC = 1.25 | | | | | | |
|---|---|---|---|---|---|---|
| **Stress** | **Sensitivity (TPR)** | **Specificity (TNR)** | **TP** | **FP** | **TN** | **FN** |
| Tetracycline | 0.88 | 0.86 | 7 | 1 | 38 | 6 |
| Kanamycin | 0.75 | 0.84 | 6 | 2 | 37 | 7 |
| Oxidative | 1.00 | 0.83 | 5 | 0 | 39 | 8 |
| Iron | 1.00 | 0.85 | 6 | 0 | 39 | 7 |

| Threshold LFC = 1.3 | | | | | | |
|---|---|---|---|---|---|---|
| **Stress** | **Sensitivity (TPR)** | **Specificity (TNR)** | **TP** | **FP** | **TN** | **FN** |
| Tetracycline | 1.00 | 0.76 | 1 | 0 | 39 | 12 |
| Kanamycin | 1.00 | 0.80 | 3 | 0 | 39 | 10 |
| Oxidative | 1.00 | 0.80 | 3 | 0 | 39 | 10 |
| Iron | 1.00 | 0.78 | 2 | 0 | 39 | 11 |

### REFERENCES


Anzai T, Imamura S, Ishihama A & Shimada T (2020) Expanded roles of pyruvate-sensing PdhR in transcription regulation of the Escherichia coli K-12 genome: fatty acid catabolism and cell motility. *Microb Genom* 6

Brosse A, Boudry P, Walburger A, Magalon A & Guillier M (2022) Synthesis of the NarP response regulator of nitrate respiration in Escherichia coli is regulated at multiple levels by Hfq and small RNAs. *Nucleic Acids Res* 50: 6753–6768

Cai S-S, Zhang L-Q, Zhang Q, Ye B-C & Zhou Y (2022) Acetylation of NarL K188 and K192 is involved in regulating Escherichia coli anaerobic nitrate respiration. *Appl Microbiol Biotechnol* 106: 7209–7221

Chen Z, Lewis KA, Shultzaberger RK, Lyakhov IG, Zheng M, Doan B, Storz G & Schneider TD (2007) Discovery of Fur binding site clusters in Escherichia coli by information theory models. *Nucleic Acids Res* 35: 6762–6777

D'Ari R (1985) The SOS system. *Biochimie* 67: 343–347

Dudin O, Geiselmann J, Ogasawara H, Ishihama A & Lacour S (2014) Repression of flagellar genes in exponential phase by CsgD and CpxR, two crucial modulators of Escherichia coli biofilm formation. *J Bacteriol* 196: 707–715

Gauger EJ, Leatham MP, Mercado-Lubo R, Laux DC, Conway T & Cohen PS (2007) Role of motility and the flhDC Operon in Escherichia coli MG1655 colonization of the mouse intestine. *Infect Immun* 75: 3315–3324

Grainger DC, Hurd D, Harrison M, Holdstock J & Busby SJW (2005) Studies of the distribution of Escherichia coli cAMP-receptor protein and RNA polymerase along the E. coli chromosome. *Proc Natl Acad Sci U S A* 102: 17693–17698

Groisman EA (2001) The pleiotropic two-component regulatory system PhoP-PhoQ. *J Bacteriol* 183: 1835–1842

Kelly A, Goldberg MD, Carroll RK, Danino V, Hinton JCD & Dorman CJ (2004) A global role for Fis in the transcriptional control of metabolism and type III secretion in Salmonella enterica serovar Typhimurium. *Microbiology* 150: 2037–2053

Landgraf JR, Wu J & Calvo JM (1996) Effects of nutrition and growth rate on Lrp levels in Escherichia coli. *J Bacteriol* 178: 6930–6936

Li Z & Demple B (1994) SoxS, an activator of superoxide stress genes in Escherichia coli. Purification and interaction with DNA. *J Biol Chem* 269: 18371–18377

Marzan LW & Shimizu K (2011) Metabolic regulation of Escherichia coli and its phoB and phoR genes knockout mutants under phosphate and nitrogen limitations as well as at acidic condition. *Microb Cell Fact* 10: 39

Partridge JD, Bodenmiller DM, Humphrys MS & Spiro S (2009) NsrR targets in the Escherichia coli genome: new insights into DNA sequence requirements for binding and a role for NsrR in the regulation of motility. *Mol Microbiol* 73: 680–694

Shimizu K (2013) Regulation Systems of Bacteria such as Escherichia coli in Response to Nutrient Limitation and Environmental Stresses. *Metabolites* 4: 1–35

Sulavik MC, Gambino LF & Miller PF (1995) The MarR repressor of the multiple antibiotic resistance (mar) operon in Escherichia coli: prototypic member of a family of bacterial regulatory proteins involved in sensing phenolic compounds. *Mol Med* 1: 436–446

Taniguchi Y, Choi PJ, Li G-W, Chen H, Babu M, Hearn J, Emili A & Xie XS (2010) Quantifying E. coli Proteome and Transcriptome with Single-Molecule Sensitivity in Single Cells. *Science* 329: 533–538

Tierrafría VH, Rioualen C, Salgado H, Lara P, Gama-Castro S, Lally P, Gómez-Romero L, Peña-Loredo P, López-Almazo AG, Alarcón-Carranza G, *et al* (2022) RegulonDB 11.0: Comprehensive high-throughput datasets on transcriptional regulation in Escherichia coli K-12. *Microb Genom* 8

Ueguchi C & Mizuno T (1993) The Escherichia coli nucleoid protein H-NS functions directly as a transcriptional repressor. *EMBO J* 12: 1039–1046

Unden G & Schirawski J (1997) The oxygen-responsive transcriptional regulator FNR of Escherichia coli: the search for signals and reactions. *Mol Microbiol* 25: 205–210

Zaslaver A, Bren A, Ronen M, Itzkovitz S, Kikoin I, Shavit S, Liebermeister W, Surette MG & Alon U (2006) A comprehensive library of fluorescent transcriptional reporters for Escherichia coli. *Nat Methods* 3: 623–628
